# Effects of transient, persistent, and resurgent sodium currents on excitability and spike regularity in vestibular ganglion neurons

**DOI:** 10.1101/2023.11.28.569044

**Authors:** Selina Baeza-Loya, Ruth Anne Eatock

## Abstract

Vestibular afferent neurons occur as two populations with differences in spike timing regularity that are independent of rate. The more excitable regular afferents have lower current thresholds and sustained spiking responses to injected currents, while irregular afferent neurons have higher thresholds and transient responses. Differences in expression of low-voltage-activated potassium (K_LV_) channels are emphasized in models of spiking regularity and excitability in these neurons, leaving open the potential contributions of the voltage-gated sodium (Na_V_) channels responsible for the spike upstroke. We investigated the impact of different Na_V_ current modes (transient, persistent, and resurgent) with whole-cell patch clamp experiments in mouse vestibular ganglion neurons (VGNs), the cultured and dissociated cell bodies of afferents. All VGNs had transient Na_V_ current, many had a small persistent (non-inactivating) Na_V_ current, and a few had resurgent current, which flows after the spike peak when Na_V_ channels that were blocked are unblocked. Na_V_1.6 channels conducted most or all of each Na_V_ current mode, and a Na_V_1.6-selective blocker decreased spike rate and altered spike waveforms in both sustained and transient VGNs. A Na_V_ channel agonist enhanced persistent current and increased spike rate and regularity. We hypothesized that persistent and resurgent currents have different effects on sustained (regular) VGNs vs. transient (irregular) VGNs. Lacking blockers specific for the different current modes, we used modeling to isolate their effects on spiking of simulated transient and sustained VGNs, driven by simulated current steps and noisy trains of simulated EPSCs. In all simulated neurons, increasing transient Na_V_ current increased spike rate and rate-independent regularity. In simulated sustained VGNs, adding persistent current increased both rate and rate-independent regularity, while adding resurgent current had limited impact. In transient VGNs, adding persistent current had little impact, while adding resurgent current increased both rate and rate-independent irregularity by enhancing sensitivity to synaptic noise. These experiments show that the small Na_V_ current modes may enhance the differentiation of afferent populations, with persistent currents selectively making regular afferents more regular and resurgent currents selectively making irregular afferents less regular.

## 1 Introduction

Vestibular hair cells transmit information about head motions and tilt to the peripheral terminals of bipolar vestibular ganglion neurons (VGNs, Fig. 1A). VGNs in amniotes are well-known for their differences in regularity of spike timing: regular and irregular VGNs synapse on hair cells in peripheral and central zones of vestibular inner ear epithelia and have tonic and phasic response dynamics, respectively (reviewed in Goldberg, 2000; Eatock and Songer, 2011). The difference in regularity is rate-independent, such that it holds even when afferents have similar average rates. Evidence from *in vivo* neuronal responses to head motions links regularity to encoding strategy: more “rate-based” in regular afferents and more “temporal” in irregular afferents, with each strategy favoring different kinds of sensory information (reviewed in Cullen, 2019). The cellular mechanisms that give rise to the difference in regularity are therefore significant for vestibular information processing. Here we examine whether the existence of different modes – transient, persistent, and resurgent – of voltage-gated sodium current (I-Na_V_) contributes to differences in afferent excitability (tendency to spike) and rate-independent regularity.

**Figure 1.**
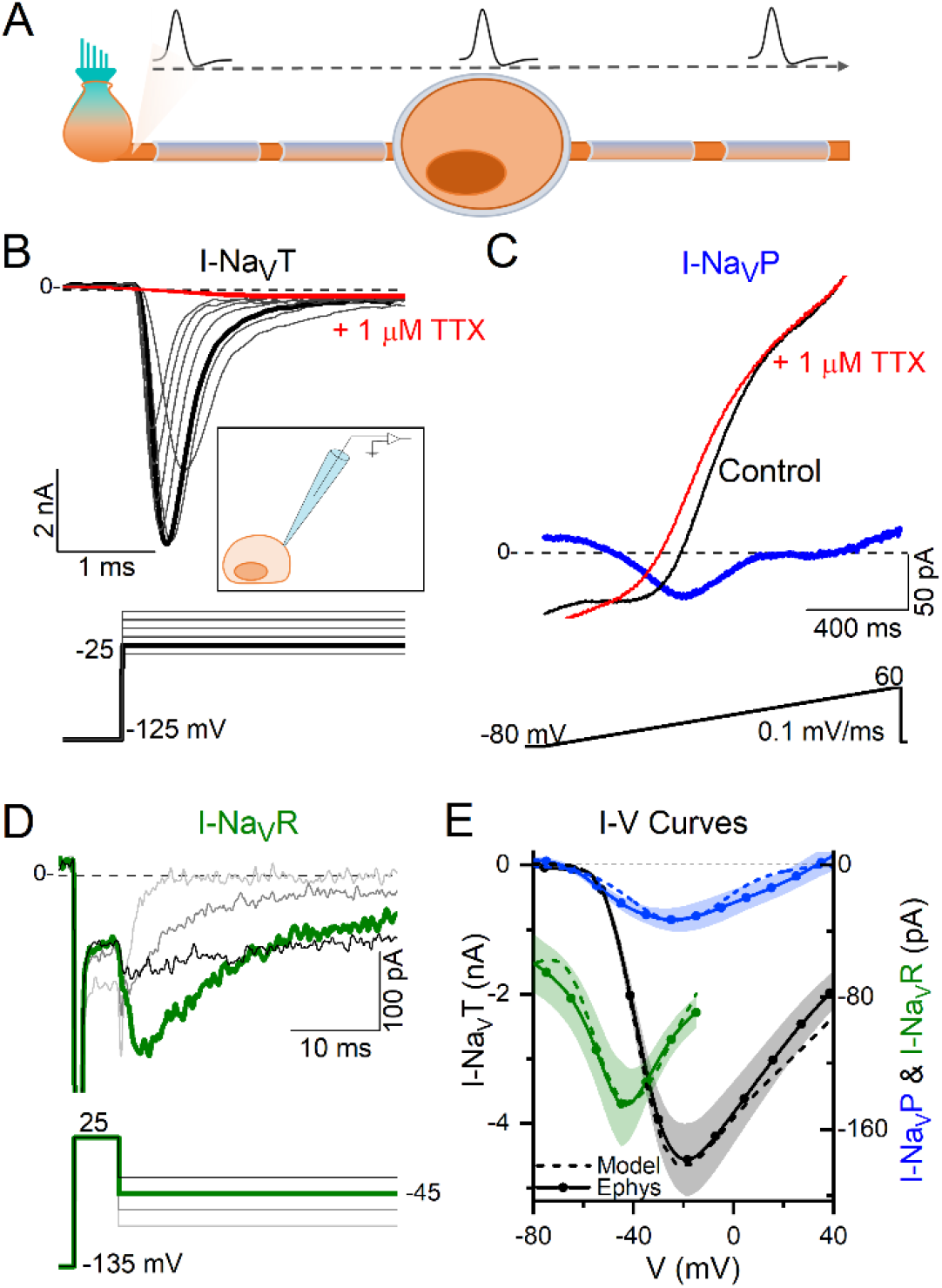
VGNs expressed Na_V_T current, many expressed Na_V_P current, and a few expressed Na_V_R currents. **(A)** VGNs are the isolated cell bodies of bipolar vestibular afferents (orange) synapsing on a hair cell (blue). *In vivo*, APs initiate at a heminode close to the synapse and adjacent to the first myelinated internode and propagate through the myelinated cell body, toward the brain. **(B)** Transient sodium current (I-Na_V_T) in a P13 VGN, evoked by stepping up from a prepulse of –125 mV in 5 mV increments. I-Na_V_T was always fully blocked by 1 μM TTX. *Inset:* Diagram of patching an isolated, demyelinated VGN. **(C)** A small, non-inactivating (persistent) current (I-Na_V_P) was isolated from a P3 VGN by applying a 0.1 mV/ms voltage ramp (–80 mV to +60 mV) and subtracting current in 1 μM TTX from control current. **(D)** Resurgent sodium current (I-Na_V_R) in a P18 VGN, evoked by applying a +25 mV prepulse followed by repolarizing steps to –25 to –80 mV; and subtracting current in 1 μM TTX from control current. **(E)** I-V curves compared for Na_V_T, Na_V_P and Na_V_R currents. Solid lines (means) plus shading (± SEM) are averaged from recordings; dashed lines show modeling of Na_V_T, Na_V_P and Na_V_R currents (see Methods).

Depolarizing current steps evoke a spectrum of firing patterns in isolated VGN cell bodies, ranging from sustained spiking to transient. Several lines of evidence indicate that sustained and transient firing patterns correspond, respectively, to regular and irregular spike timing *in vivo* (Iwasaki et al., 2008; Kalluri et al., 2010; Ono et al., 2020). *In vivo*, spikes initiate in the peripheral neurite at the heminode near its hair cell synapse and travel through the cell body into the central neurite (Fig. 1A). Over the first postnatal week, the afferent nerve fibers, including the cell body, become myelinated. We investigated the Na_V_ currents that give rise to firing patterns in dissociated VGN cell bodies. Their compact form allows the high-quality voltage clamp necessary for recording very fast Na_V_ currents. To gain access for the patch electrode to the VGN membrane beyond the first postnatal week, we cultured dissociated cell bodies overnight, which loosens or eliminates the myelin covering.

With this approach, previous work on rat VGNs in the first two postnatal weeks established that, relative to sustained VGNs, transient VGNs are less excitable, with higher current thresholds for spiking, reflecting their more negative resting potentials and lower input resistances (Iwasaki et al., 2008; Kalluri et al., 2010). A major factor in these differences is the greater expression of low-voltage-activated potassium (K_LV_) channels from the K_V_1 and K_V_7 channel families in transient VGNs. By stimulating VGNs with trains of simulated excitatory postsynaptic currents (EPSCs) with pseudo-random timing, Kalluri et al. (2010) showed that transient VGNs have larger K_LV_ currents that drive irregular spike timing. Whether diversity in Na_V_ currents contributes to afferent differences in regularity and excitability has not been established.

Na_V_ currents drive the rising phase of most neuronal action potentials (reviewed in Bean, 2007). In rat vestibular ganglia (Liu et al. 2016), we showed diverse expression of Na_V_ channel subunits and characterized multiple transient Na_V_ currents (I-Na_V_T) in immature VGNs. Here we investigated a different aspect of Na_V_ current expression in VGNs that becomes a factor as the inner ear matures: the expression of persistent (Na_V_P) or resurgent (Na_V_R) Na_V_ modes of current, which flow through the same channel subunits that produce larger Na_V_T currents (Brown et al., 1994; Raman and Bean, 1997; Meredith and Rennie, 2020).

Most Na_V_ channels strongly inactivate within milliseconds after depolarization, producing I-Na_V_T. In some cases, however, a small fraction do not inactivate even after seconds, creating I-Na_V_P. I-Na_V_P activates more negatively than I-Na_V_T and therefore enhances Na_V_ channel availability near resting potential, contributing to excitability and repetitive firing (Crill, 1996; Raman & Bean, 1997; Do and Bean, 2003). In another type of Na_V_ current, Na_V_ channels opened by a strong depolarization (like a large spike) are blocked by an intracellular molecule before they inactivate; upon repolarization, the channels rapidly unblock and carry I-Na_V_R until they deactivate (Raman & Bean, 1997; Aman & Raman, 2024). These channels enhance depolarizing current on the downstroke of a spike and during the interspike interval.

Na_V_P and Na_V_R current modes have been recorded from gerbil calyces, the large terminals formed by vestibular afferents on type I hair cells and are attributed to tetrodotoxin (TTX)-sensitive Na_V_1.6 channels (Meredith and Rennie, 2020). We speculated that diversity in Na_V_ current modes contributes to differences in excitability and/or regularity in VGNs, as suggested for certain brain neurons (Lewis and Raman, 2014). Our experiments and simulations together suggest that increasing I-Na_V_T drives increases in spike rate and regularity in all VGNs, but that persistent and resurgent currents may enhance differentiation of spike regularity between sustained (regular) and transient (irregular) vestibular afferent neurons.

## 2 Methods

### 2.1 Electrophysiology

#### Animals

For all electrophysiology experiments, wildtype CD-1 mice were obtained from Charles River Laboratories (Wilmington, MA). Mice were housed at the University of Chicago Animal Care Facility and were handled in accordance with the National Research Council Guide for the Care and Use of Laboratory Animals. All procedures were approved by the animal care committee at the University of Chicago.

#### Preparation

Whole-cell currents and voltages were recorded from VGNs isolated from mice on postnatal day 3 (P3) to P25 (11 ± 0.5 (SEM) days old, median = 8, n = 146). VGNs from mice of both sexes were pooled into cell cultures; in the few cases where cells were from cultures made of a single sex (n = 7 female, 8 male cells), there were no significant differences in Na_V_ current voltage dependence of activation (p = 0.99, power = 0.05) or conductance density (p = 0.18, power = 0.27) between sexes.

Mice were anesthetized via isoflurane inhalation and decapitated. Temporal bones were dissected in chilled Liebovitz-15 (L-15) medium supplemented with 10 mM HEPES to pH 7.4, ∼320 mmol/kg. Chemicals were purchased from ThermoFisher (Waltham, MA) unless otherwise stated.

Each otic capsule was exposed, and the superior compartment of the vestibular ganglion was dissected out. The superior ganglion houses the cell bodies of VGNs that synapse on hair cells in the utricle, part of the saccule, and the lateral and anterior semicircular canals, and project centrally to the vestibular nuclei in brain stem and cerebellum. We followed a dissociation protocol as described in Limón et al., 2005, with small changes. The tissue was placed in L-15 supplemented with 0.12% collagenase and 0.12% trypsin for 15-20 minutes at 37°C. The ganglion was then dissociated by gentle trituration into Minimal Essential Medium with Glutamax supplemented with 10 mM HEPES, 5% fetal bovine serum, and 1% penicillin (Sigma-Aldrich, St. Louis, MO). Cells were allowed to settle in glass-bottomed culture dishes (MatTek, Ashland, MA) pre-coated with poly-D-lysine. In most experiments, recordings were made after cells were incubated 10-20 hours in 5% CO_2_ - 95% air at 37°C. Overnight incubation reduced myelin and satellite cell coverage from cell bodies. Age of cells does not include time in culture: e.g., data from a P17 VGN indicates the pup was 17 days old when neurons were harvested, and the cells were P17 + ∼1 day *in vitro*. The number of surviving cells decreased with age of animal.

#### Recording solutions

During experiments, cells were bathed in either one of two external solutions, summarized in Table 1. For voltage clamp experiments, we used a “Na^+^-reduced” external solution and a “Cs^+^” -based internal solution, designed to minimize K^+^ currents, Ca^2+^ currents, and reduce Na^+^ currents. K^+^ currents were minimized and Na^+^ currents reduced by substituting Cs^+^ for K^+^ (in both external and internal solutions, Table 1) and replacing about half of the external Na+ with equivalent tetraethylammonium chloride (TEACl, a K channel blocker). By reducing K^+^ and Na_V_ current, these solutions improved voltage clamp quality by reducing voltage errors and the impact of the clamp time constant. To minimize Ca^2+^ current, only trace Ca^2+^ was present and Mg^2+^ was added to block voltage-gated calcium (Ca_V_) channels. For current clamp experiments to gather spiking data, we used more physiological external and internal solutions (“standard” solutions, Table 1).

**Table 1.** Whole-cell recording solutions. pH was adjusted with CsOH for Na^+^ reduced external and Cs internal solutions, NaOH for standard external solution, and KOH for standard internal solution.

| Table 1: Whole-cell recording solutions |  |  |  |  |  |  |  |  |  |  |  |  |
| --- | --- | --- | --- | --- | --- | --- | --- | --- | --- | --- | --- | --- |
| External |  |  |  |  |  |  |  |  |  |  |  |  |
| in mM | NaCl | KCl | CsCl | $\text{CaCl}_2$ | $\text{MgCl}_2$ | TEACl | $\text{NaH}_2\text{PO}_4$ | D-Glucose | | HEPES | pH | Osm (mmol/kg) |
| $\text{Na}^+$ Reduced | 75 | 0 | 5.4 | 0 | 2.5 | 75 | 0 | 10 | | 5 | 7.4 | 310 |
| Standard | 144 | 5.1 | 0 | 1.3 | 0.9 | 0 | 0.7 | 5.6 |  | 10 | 7.4 | 310 |
| Internal |  |  |  |  |  |  |  |  |  |  |  |  |
| in mM | KCl | CsCl | $\text{CaCl}_2$ | $\text{MgCl}_2$ | $\text{Na}_2$ creatine phosphate | Mg-ATP | Li-GTP | Na-cAMP | EGTA | HEPES | pH | Osm (mmol/kg) |
| $\text{Cs}^+$ | 0 | 148 | 0.8 | 0 | 3.5 | 3.5 | 0.1 | 0.1 | 5 | 5 | 7.4 | 300 |
| Standard | 135 | 0 | 0.1 | 0.5 | 5 | 3 | 0.1 | 0.1 | 5 | 5 | 7.25 | 285 |

#### Whole cell recordings

Cells were visualized at 600X on an inverted microscope equipped with Nomarski optics (IMT-2; Olympus, Lake Success, NY). We chose round cells with diameters >5 µm (range 8 – 25 µm). Mean membrane capacitance measured by the amplifier was 15 ± 0.5 (median = 16, n = 135). No correlation between age and cell size was observed. Neurons were distinguished from glia by their shape and the presence of Na_V_ currents and/or spikes.

Stimuli were delivered, and responses recorded and amplified, with a Multiclamp 700B amplifier, Digidata 1440A digitizer, and pClamp 11 software (Axon Instruments, Molecular Devices, Sunnyvale, CA), with low-pass Bessel filtering set at 10 kHz and sampling interval set at 5 µs. Stimuli include voltage steps and ramps, current steps, and synthetically generated frozen trains of excitatory postsynaptic currents (EPSCs) based on recordings from vestibular terminals, as in Kalluri et al., 2010. Electrodes were pulled from soda glass (King Precision Glass, Claremont, CA) to resistances of 3 – 4 MΩ in our solutions and wrapped with parafilm to reduce pipette capacitance. All membrane voltages were corrected offline for a liquid junction potential of either 4.7 (standard solution) or 5.2 mV (reduced solution), calculated with JPCalc software (Barry 1994) in pClamp 11.

The series resistance (R_S_) ranged between 3 and 10 MΩ and was compensated by 73 ± 0.9% (n = 134). Recordings were analyzed only when the seal exceeded 1 GΩ and, if standard solutions were used, the cell maintained a stable resting membrane potential more negative than −60 mV. After compensation for junction potentials, holding potential in voltage-clamp mode was −65 mV (−64.7 mV in standard solutions, −65.2 mV in Na^+^ Reduced/Cs^+^ solutions). Input resistance measured at holding potential was 712 ± 56 MΩ. Fast Na_V_ currents are difficult to voltage clamp at body temperature, so we recorded at room temperature (23 – 25°C). For our average cell capacitance (15 pF) and after R_S_ compensation, the voltage clamp time constant ranged from ∼15-50 *µ*s, adequate for recording fast Na_V_ currents at room temperature, at which time-to-peak and inactivation time constants are ∼1 ms, and activation and deactivation time constants are shorter (Patel et al. 2015; Alexander et al., 2021).

#### Pharmacology

On the day of experiments, we thawed stock solutions of pharmacological agents: Na_V_ channel blockers tetrodotoxin (TTX; Enzo Life Sciences, Farmingdale, NY; 2 mM in distilled water) and 4,9-anhydro-tetrodotoxin (4,9-ah-TTX; Alomone Labs, Jerusalem, Israel; 500 μM in methanol), or Na_V_ channel agonist *Anemonia viridis* toxin 2 (ATX-II; Alomone Labs, Jerusalem, Israel; 100 μM in distilled water). Aliquots were added to 5 mL of external solution for final concentrations of 1 μM TTX, 100 nM 4,9-ah-TTX, and 100 nM ATX-II, chosen to maximize blocking or agonizing effects (Oliveira et al., 2004; Rosker et al., 2007). 1 mg/ml of bovine serum albumin was added to the ATX-II solution to reduce adhesion to the plastic delivery tubing. Toxin-containing solutions were applied via local perfusion (Perfusion Fast-Step, Warner Instruments, Holliston, MA) delivered with a Bee Hive Controller (BASI, West Lafayette, IN). Control and drug solutions flowed through adjacent delivery tubes and a stepper mechanism selected the tube directed at the patched cell. This system allows for no dead volume. Perfusion of control solution at the beginning of each drug series provided additional control for flow effects.

#### Analysis and statistical treatments

Analysis was performed in Clampfit (pClamp 10 or 11, Molecular Devices) and MATLAB 2023b (The MathWorks, Natick, MA). Statistical analyses, curve fitting, and figures were done in Origin Pro 2018 or 2024 (OriginLab, Northampton, MA). Means ± SEM are presented. In box plots, box indicates SEM, middle line indicates median, open squares indicate the mean, and whiskers mark 5-95% confidence intervals. For comparisons between 2 groups, we tested for normal distributions and homogeneity of variance (Levene’s test), and estimated statistical significance with Student’s t-test or, if variances were unequal, Welch’s t-test. We used paired t-tests for drug effects on individual cells and an alpha level of 0.05 for all statistical tests. For significant results, we calculated effect size with bias-corrected Hedges’ g (small effect = 0.2, medium = 0.5, large = 0.8; Durlak, 2009). To compare >2 groups, we used one-way ANOVA followed by the Bonferroni test for multiple comparisons.

The voltage dependencies of current activation and inactivation were analyzed for currents collected with R_S_ voltage error <10 mV at peak current. Activation curves of conductance (*G*) *vs*. voltage (*V*) were generated by dividing peak current (*I*) by the driving force (*V* − *E*_*rev*_) to obtain *G. E*_*rev*_ was approximated by the equilibrium potential for Na^+^ (E_Na_) for the specified Na^+^ concentrations (Table 1). The resulting *G-V* curves were generally well described by fitting a simple Boltzmann function using the Levenberg-Marquardt fitting algorithm as implemented in OriginPro:

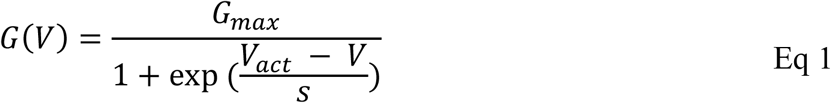

where *G*_*max*_ is the maximum conductance, *V*_*act*_ is the voltage of half-maximal activation, and *s* is the slope factor.

Inactivation *G-V* curves were generated by measuring how iterated prepulse voltages affect the conductance evoked by a test pulse (step) in the activation range (–35 or –15 mV). Peak current evoked by the test pulse was divided by driving force, plotted against the iterated prepulse voltage, and fit by a Boltzmann function:

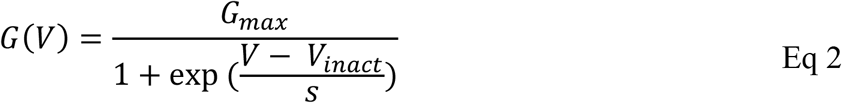

with *V*_*inact*_ equaling the voltage of half-maximal inactivation.

Coefficient of variation (*CV*) was used to measure regularity of spike trains and was calculated for a given spike train from the mean and standard deviation of interspike interval (*ISI*):

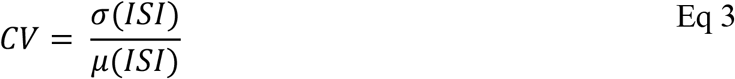

To compare intrinsic, rate-independent, regularity, we wished to compare CV at similar rates. Because CV varies inversely with spike rate, we manipulated the input current amplitude to hold spike rate approximately constant between control and experimental conditions. Events were counted as spikes when they crossed a voltage threshold, between –20 and 0 mV, which was set for each neuron to exclude subthreshold excitatory postsynaptic potentials (EPSPs). We confirmed that APs and not EPSPs were counted by comparing the waveforms: APs had higher peak values, faster rise times and decays, and, often, afterhyperpolarizations (AHPs, measured as the difference between the resting potential and the most negative potential following a spike).

### 2.2 Modeling

A Hodgkin-Huxley model of neuronal firing was used to assess the individual impacts of I-Na_V_T, I-Na_V_R and I-Na_V_P on spiking. The single-compartment model was implemented in MATLAB 2023b as a differential equation in which the net current across the neuronal membrane was taken as the sum of various individual currents (Hodgkin and Huxley, 1952):

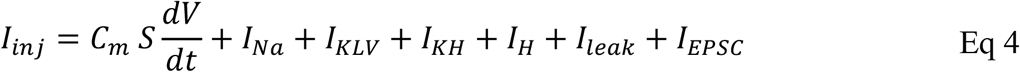

This model is an extension of the single-compartment VGN model developed by Hight and Kalluri (2016) and Ventura and Kalluri (2019) to fit rat VGN data. Membrane voltage *V(t)* was solved numerically using a backwards difference method. The specific membrane capacitance (*C*_*m*_) was fixed at 0.9 μF/cm^2^ (Gentet et al., 2000). Cell surface area (*S*) was fixed to yield a net capacitance of 15 pF, the average for our recorded mouse VGNs. The 5 ionic currents represent key current types in vestibular ganglion neurons: voltage-gated sodium (*I*_*Na*_), low-voltage-activated potassium (*I*_*KLV*_), high-voltage-activated potassium (*I*_*KH*_), hyperpolarization-activated cyclic nucleotide-gated (*I*_*H*_), and leak (*I*_*leak*_). The model VGN was stimulated either by injected current steps (*I*_*inj*_) with *I*_*EPSC*_ set to 0, or with trains of simulated vestibular excitatory postsynaptic currents (pseudo-EPSCs, *I*_*EPSC*_) with *I*_*inj*_ set to 0. A holding period of 500 ms preceded any current injection to allow the resting membrane potential to equilibrate.

To ensure that the combinations of parameters for currents reproduced APs and firing patterns of different VGNs observed in whole-cell recordings (see Fig. 5A and C), we fit the model output to recorded data using a local search optimization algorithm. This algorithm compared the phase plane plots of model APs, produced by different combinations of ionic, leak, and injected currents, against the phase plane plots of averaged recorded APs (Fig. S1, Table 2). We constrained the Na_V_ conductance parameter range to the specific range recorded for each neuron type in this study (e.g., Fig 5B) and, for the other ionic conductances, determined the ranges based on previously published work (see Table 2 for citations). For each firing pattern, the combination of parameters yielding a simulated AP with the lowest mean squared error relative to the average recorded AP was accepted; the resulting simulations are shown in Fig. S1: panel D (AP waveforms), E (phase plane plots) and F (firing patterns). The model parameter combinations for the 4 identified VGN firing patterns (Fig. S1C, F) are summarized in Table 2.

**Table 2.** Conductance parameters used for spiking by model VGNs. Sources: ^a^Hight and Kalluri (2016), ^b^Kalluri et al. (2010), ^c^Iwasaki et al. (2008), ^d^Ventura and Kalluri (2019). g_NaVT_ and V_rest_ from this study.

| Table 2: Conductance parameters used for spiking |  |  |  |  |  |  |
| --- | --- | --- | --- | --- | --- | --- |
| Firing Pattern | $\bar{g}_{\text{NaVT}}$ (mS/cm<sup>2</sup>) | $\bar{g}_{\text{KLV}}$ <sup>a,b,d</sup> | $\bar{g}_{\text{KH}}$ <sup>a,c,d</sup> | $\bar{g}_{\text{H}}$ <sup>d</sup> | $\bar{g}_{\text{leak}}$ <sup>a,d</sup> | $V_{\text{rest}}$ (mV) |
| Sustained-A | 16 | 0 | 4.5 | 0.2 | 0.02 | -60.1 |
| Sustained-B | 13 | 0.2 | 4 | 0.5 | 0.05 | -63.5 |
| Sustained-C | 11 | 0.5 | 4 | 0.1 | 0.05 | -64.1 |
| Transient | 7 | 1.2 | 2.5 | 0.9 | 0.1 | -65.7 |

In previous versions of the model (Hight and Kalluri, 2016; Ventura and Kalluri, 2019), *I*_*Na*_ was entirely transient (I-Na_V_T) and based on the formulation in Rothman and Manis (2003). Here, we adapted *I*_*Na*_ to include persistent (I-Na_V_P) and resurgent (I-Na_V_R) Na_V_ currents. We used I-Na_V_T equations from Hight and Kalluri (2016) and I-Na_V_R and I-Na_V_P equations from Venugopal et al. (2019) and Wu et al. (2005), substituting in our mouse VGN values for conductance density and voltage dependence. Figure 1E shows the *I-V* relations of both recorded (continuous lines) and modeled (dashed lines) Na_V_ current components.

The equation for total Na_V_ current was based on the computational model by Venugopal et al. (2019) and can be written as:

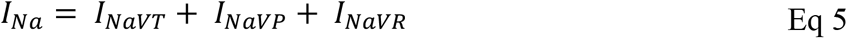

where *I*_*NaVT*_, *I*_*NaVP*_, and *I*_*NaVR*_ are modeled as:

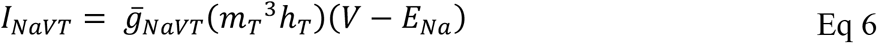

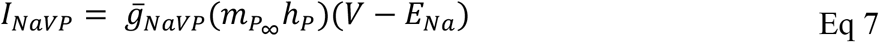

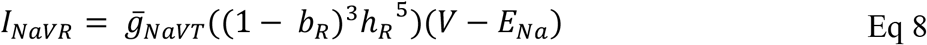

Model parameters for Na_V_ current modes are summarized in Table 3. Conductance densities 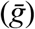 cover the range of experimentally derived values from this study on VGN cell bodies and the data of Meredith and Rennie (2020) on Na_V_ currents in VGN calyx terminals on hair cells: maximal persistent conductance 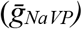 was set to 3% of the transient conductance 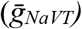 and the maximum resurgent conductance 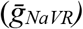 was set to 10% of 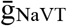, as recorded from cell bodies in this study and as recorded in vestibular afferent (calyceal) terminals (Meredith and Rennie, 2020).

**Table 3.**
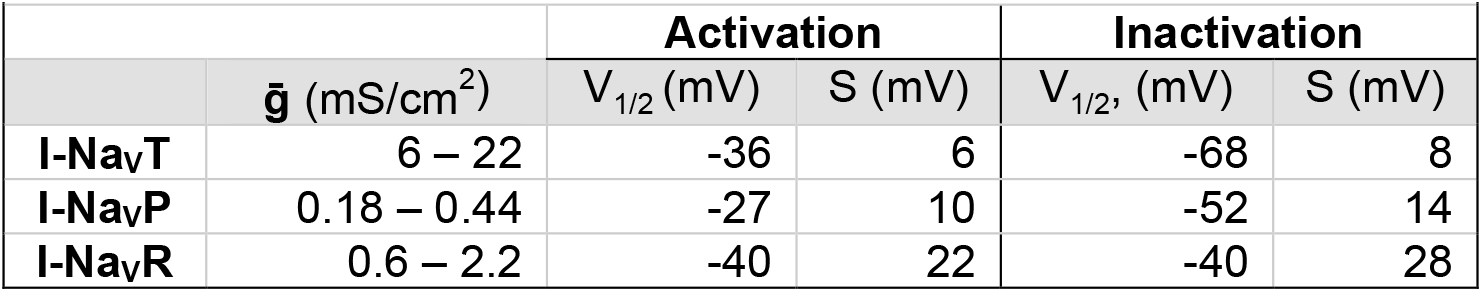
Steady-state activation and inactivation parameters used in modeling Na_V_ current modes.

#### Transient sodium current

The steady-state voltage-dependent activation (m_T_) and inactivation (h_T_) of Na_V_T currents are modeled as in Rothman and Manis (2003), with voltage of half-activation (*V*_*1/2*_) and slope factor (*s*) set equal to the mean values from this study.

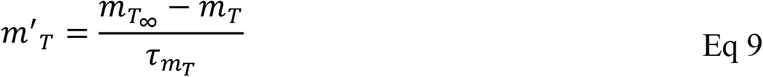

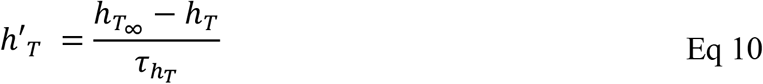

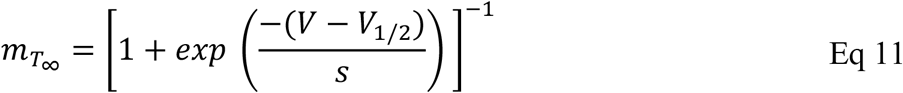

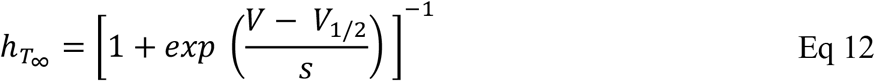

Steady state voltage-dependent time constants of activation and inactivation functions for Na_V_T currents, originally from Rothman and Manis (2003), remained mostly unchanged:

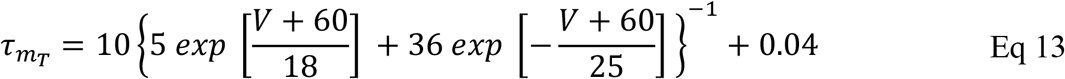

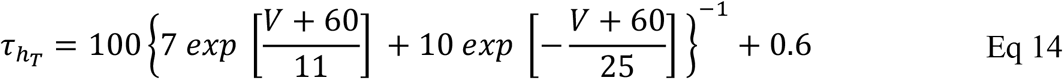

#### Persistent and resurgent sodium currents

Steady-state activation (h_P_), inactivation (m_P_), and voltage-dependent time constant of inactivation (τ_hP_) for Na_V_P current are based on Venugopal et al. (2019) and Wu et al. (2005) and are modeled as:

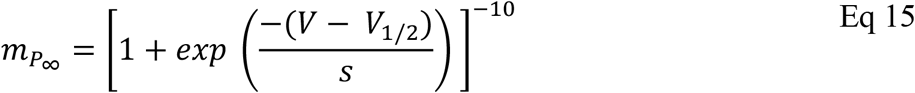

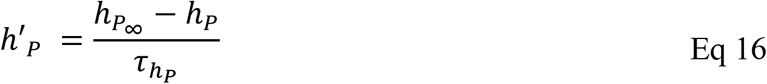

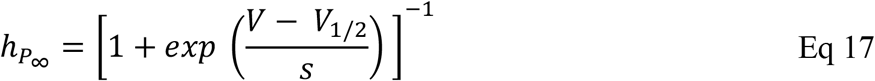

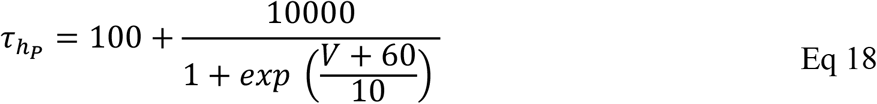

The formulation for I-Na_V_R is from Venugopal et al. (2019). It alters the Hodgkin-Huxley conductance-based formulation to incorporate state-dependent Na^+^ resurgence due to unblocking of a channel that was blocked upon opening. The equations that govern voltage-dependent blocking/unblocking (*b*_*R*_) kinetics are as follows:

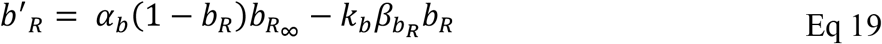

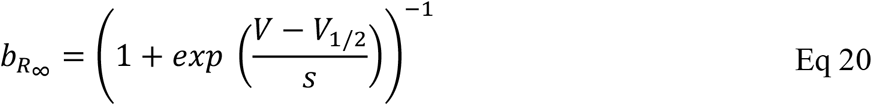

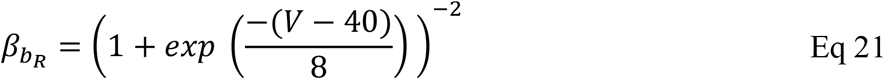

where constants *α*_*b*_ (0.08) and *k*_*b*_ (0.9) control the rate of unblocking. The voltage-dependent inactivation of I-Na_V_R (*h*_*R*_) functions include:

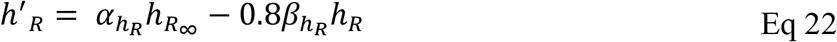

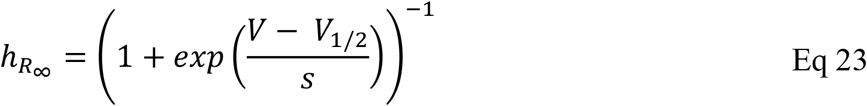

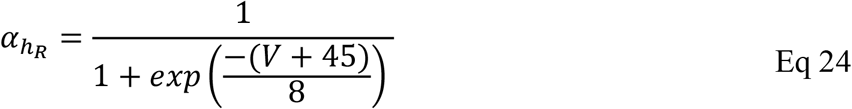

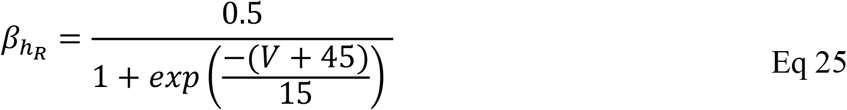

#### Synaptic conductance and EPSC shape

Synaptic input was generated and modeled as described in Hight and Kalluri (2016). Briefly, modeled synaptic events were randomly drawn from Gaussian distributions of size and timing based on EPSC amplitudes and rates. For our model simulations, we simulated EPSCs with a shape based on EPSCs of vestibular afferent calyces in the excised P8 CD-1 mouse utricular epithelium (lateral extrastriolar zone) at room temperature. These have longer onset and decay times than the S1 EPSC shape used in Hight and Kalluri (2016) (Fig. S2). An exponential function was fitted to an averaged synaptic event:

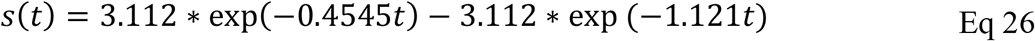

## 3 Results

Whole-cell ruptured-patch recordings were taken from somata of vestibular ganglion neurons (VGNs; n = 146) dissociated from CD1 mice of both sexes between P3 and P25 (median = P8) and cultured overnight. VGN cell bodies allow better space clamp than afferents recorded *in vivo*.

Because Na_V_P and Na_V_R current modes upregulate with development (Browne et al., 2017; Hong et al., 2018; Meredith and Rennie, 2020), we needed to record beyond the first postnatal week, by which time myelin is extensive. With overnight culture, VGN cell bodies lose their myelin wrapping, allowing access for recording, but have not yet acquired long neurites that occur with long term culture.

First, we show Na_V_T, Na_V_P, and Na_V_R current modes recorded from VGNs, and their contributions to voltage dependence and time course of the overall Na_V_ current (I-Na_V_). We describe experiments investigating contributions of a channel subunit, Na_V_1.6, that carries all three current modes. We then assess the roles of Na_V_ currents in AP waveforms and spiking during current clamp. Finally, we examine the roles of each current mode in a computational model of VGN spiking.

### 3.1 Properties of Na_V_ Currents in VGNs

To collect and characterize Na_V_ currents, we used solutions that lowered R_S_ voltage errors by decreasing Na_V_ currents: “Cs^+^” internal solution and “Na^+^ Reduced” external solution (Table 1), which had Cs^+^ instead of K^+^ and no added Ca^2+^ to minimized contamination with K^+^ or Ca^2+^ current. We recorded at room temperature (∼24°C) to slow activation speed of fast Na_V_ currents into the range for which our voltage clamp time constant is adequate (∼15-50 *µ*s, see Methods).

#### All VGNs had I-Na_V_T, some had I-Na_V_P, and few had I-Na_V_R

Figure 1 shows exemplar Na_V_ currents recorded from VGNs in response to voltage protocols previously designed to reveal Na_V_T, Na_V_P, and Na_V_R current components (Stafstrom et al., 1982; Raman and Bean, 1997). Depolarizing steps following a hyperpolarizing prepulse revealed fast-inactivating I-Na_V_T in all VGNs (Fig. 1B), as previously reported (Chabbert et al., 1997; Risner and Holt, 2006; Liu et al., 2016). Table 4 summarizes I-Na_V_T properties.

**Table 4.** Electrophysiological properties of Na_V_ currents in VGNs. Asterisks indicate significance in one or two comparisons. See Table S1 for full summary of statistical analyses.

| Table 4: Electrophysiological properties of $Na_V$ currents in VGNs | | | | | | |
| --- | --- | --- | --- | --- | --- | --- |
|  |  | Activation |  | Inactivation |  | G density<br>(nS/pF) |
| | | $V_{1/2}$<br>(mV) | S<br>(mV) | $V_{1/2}$<br>(mV) | S<br>(mV) | |
| Nav current | I-$Na_V$<br>(Na<sup>+</sup> Red Ext Sol'n) | -35.8 ± 0.7<br>(n = 75) | 5.7 ± 0.2<br>(76) | -68.1 ± 1.1<br>(73) | 8.4 ± 0.5<br>(74) | 3.8 ± 0.3<br>(74) |
| | I-$Na_V$<br>(Standard Ext Sol'n) | -38.9 ± 1.2<br>(29) | 3.2 ± 0.3<br>(29) | -59.1 ± 1.6<br>(23) | 4.4 ± 0.8<br>(23) | 6.8 ± 0.7<br>(29) |
| | I-$Na_V$<br>(Liu et al., 2016, rat<br>VGN) | -36.5 ± 1.6<br>(11) | 5.7 ± 0.4<br>(11) | -76.3 ± 0.2<br>(12) | 7.6 ± 0.1<br>(12) | 8.7 ± 1.0<br>(12) |
| I-Nav modes<br>(Figure 2) | I-$Na_V$ (T) | -35.6 ± 1.0<br>(10) | 6.6 ± 0.3<br>(10) | -70.1 ± 2.1<br>(10) | 12.3 ± 1.3<br>(10) | 2.9 ± 0.3<br>(31) |
| | I-$Na_V$ (T+P) | -37.7 ± 1.6<br>(12) | 7.1 ± 0.4<br>(12) | -66.4 ± 2.3<br>(12) | 12.8 ± 0.8<br>(12) | 3.6 ± 0.4<br>(34) |
| | I-$Na_V$ (T+P+R) | -48.6 ± 5.7*<br>(6) | 6.0 ± 0.4<br>(6) | -67.5 ± 6.3<br>(6) | 11.5 ± 1.0<br>(6) | 5.9 ± 0.9*<br>(6) |
| 4,9-ah-TTX<br>(Figure 3) | Control (I-Nav) | -38.7 ± 1.3<br>(12) | 5.7 ± 0.5<br>(12) | -68.3 ± 2.6<br>(12) | 9.1 ± 1.4<br>(12) | 3.1 ± 0.4<br>(13) |
|  | Subtraction | -39.4 ± 1.6 | 4.7 ± 0.6 | -68.4 ± 2.4 | 8.8 ± 1.3 | 2.4 ± 0.3 |
|  | Residual | -42.3 ± 2.3 | 9.2 ± 2.1 | -80.6 ± 3.8* | 14.5 ± 4.9 | 0.9 ± 0.3* |
| ATX-II<br>(Figure 4) | Control (I-Nav) | -37.7 ± 1.2<br>(7) | 5.7 ± 0.3<br>(7) | -76.7 ± 3.6<br>(7) | 11.2 ± 0.8<br>(7) | 3.7 ± 1.7<br>(7) |
|  | ATX-II | -38.7 ± 1.3 | 5.8 ± 0.1 | -80.9 ± 2.9 | 16.3 ± .3* | 1.9 ± 0.4 |

To reveal Na_V_P current, we eliminated rapidly inactivating I-Na_V_T by applying a slow depolarizing ramp (0.1 mV/ms from −80 to +60 mV) and obtained the TTX-sensitive non-inactivating component by subtracting the ramp current remaining in 1 μM TTX (Fig. 1C). This method often revealed I-Na_V_P as a small TTX-sensitive inward current activating above –70 mV. I-Na_V_P was evident in 42 of 78 (54%) VGNs, P3-25, always in combination with I-Na_V_T and in 4 cases with I-Na_V_R as well.

I-Na_V_R was revealed by stepping briefly from –125 mV to +25 mV to open Na_V_ channels, then repolarizing to iterated negative voltages (Fig. 1D) (Raman and Bean, 1997). It is thought that a blocking particle enters the channel as it opens at the activating voltage and prevents inactivation; with repolarization the channel unblocks, yielding I-Na_V_R, which then inactivates (Bant and Raman, 2010; White et al., 2019). Overall, I-Na_V_R was much less common than I-Na_V_P, occurring in just 6 of 78 VGNs (8%) tested in voltage clamp. All the cells with I-Na_V_R were older than P10; for this age group, the incidence was 6/49 or 12%. Two of the 6 VGNs had I-Na_V_T and I-Na_V_R but no I-Na_V_P. Developmental upregulation of I-Na_V_R has been described in vestibular afferent endings, where I-Na_V_R was found in 35/67 (54%) calyces >P30 (Meredith and Rennie, 2020).

Relative to I-Na_V_T, which for iterated voltage steps peaked at −20 mV, I-Na_V_R and I-Na_V_P reached maximal amplitude at –45 mV repolarization voltage and −25 mV ramp voltage, respectively (Fig. 1E). Although, on average, peak I-Na_V_P and peak I-Na_V_R were just 1% and 3% of peak total I-Na_V_, the small Na_V_P and Na_V_R currents can be relatively more substantial at subthreshold voltages; for example, from −60 to −50 mV, average I-Na_V_P is 13% and I-Na_V_R is 124% of the peak I-Na_V_T over the same voltage range. Note that I-Na_V_R is present only following a depolarizing prepulse or, *in vivo*, an AP.

I-Na_V_T, I-Na_V_P, and I-Na_V_R were all completely blocked by 1 μM TTX. In contrast, in “acute” recordings from immature rat VGNs (P < 8) on the day of dissociation, Liu et al. (2016) recorded multiple kinds of I-Na_V_T with different TTX sensitivities and kinetics: TTX-insensitive Na_V_1.5 current (IC_50_ ∼300 nM TTX) and TTX-resistant Na_V_1.8 current (no block at 5 μM TTX), in addition to TTX-sensitive current. As in other studies of overnight-cultured VGNs (Chabbert et al., 1997; Risner and Holt, 2006; Liu et al. 2016), we did not detect TTX-insensitive or -resistant currents, even with 300 nM TTX to block the large TTX-sensitive currents (n = 5, not shown).

#### The addition of I-Na_V_P and I-Na_V_R affects overall activation voltage

Cells with I-Na_V_R had larger average peak current density relative to VGNs with both Na_V_T and Na_V_P currents (I-Na_V_(T+P)) or just I-Na_V_T (Fig. 2A; collected with protocol shown in Fig. 1A). The 2 VGNs with I-Na_V_(T+R) are shown separately in Figure 2A and B and had unusually large current densities (Fig. 2C, green) and negative midpoints of activation and inactivation (Fig. 2B).

**Figure 2.**
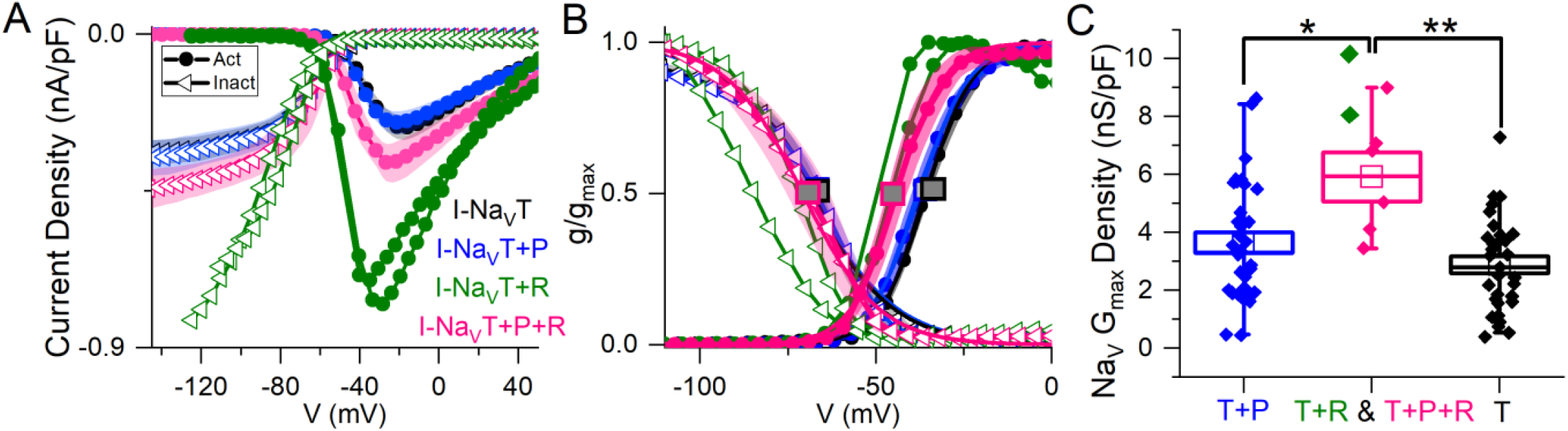
Voltage dependence of total Na_V_ current varied with different combinations of Na_V_T, Na_V_P and/or Na_V_R current modes. **(A)** I-V curves averaged for cells with different current modes. 2 VGNs with I-Na_V_T plus I-Na_V_R (green) had large current densities and are shown individually. **(B)** G-V activation curves show that voltage dependence differed in cells with different combinations of Na_V_ current modes. V_1/2_ of activation and inactivation are marked with square symbols. Voltage dependence of activation was more negative in VGNs with I-Na_V_R (I-NaV(T+R) and I-NaV(T+P+R); green and pink lines) than I-Na_V_(T+P) (p = 0.02, blue) and I-Na_V_T (p = 0.007, black). V_1/2_ of inactivation did not differ (one-way 3-factor ANOVA, p = 0.7). **(C)** VGNs with I-Na_V_R, I-Na_V_(T+P+R) (pink) and I-Na_V_(T+R) (green), had larger (pooled) conductance densities than VGNs without I-Na_V_R: I-Na_V_(T+P) (blue; one-way 3-factor ANOVA, p = 0.02) and I-Na_V_T (black; p = 0.002).

To measure how Na_V_P and Na_V_R currents influenced the voltage dependence of the total Na_V_ current, we fit activation and inactivation (peak G-V) curves with Boltzmann functions (Eqs. 1 and 2; Fig. 2B) and compared fit parameters (Table 4; statistical tests in Table S1). There was no difference detected in half-maximal activation potential (V_1/2, Act_) values between VGNs with I-Na_V_T and I-Na_V_(T+P) (Fig. 2B); although the statistical power is low, the lack of clear difference is not surprising given the small size of I-Na_V_P. Cells with I-Na_V_R (n = 6), however, had V_1/2, Act_ values shifted significantly negative to cells with I-Na_V_(T+P) or just I-Na_V_T (Fig. 2B). The negative shift of activation voltage suggests that I-Na_V_R may decrease the current threshold for spiking, possibly by reducing overall rates of inactivation. Half-maximal inactivation potential (V_1/2, Inact_) did not differ significantly across groups (Fig. 2B).

In Figure 2C, peak current density values have been converted to maximum Na_V_ conductance density (Na_V_ G_Max_ density). To assess the effect of I-Na_V_R, we pooled all VGNs with I-Na_V_R [(T+P+R) and (T+R)]. VGNs with I-Na_V_(T+P+R) had greater total Na_V_ G_Max_ density relative to I-Na_V_T and I-Na_V_(T + P) (Table 4). This indicates that VGNs with resurgent current have a greater I-Na_V_ conductance. Later (see Fig. 8), we use the computational model to compare the effects of increasing total conductance with just I-Na_V_T current vs. I-Na_V_T plus P and/or R modes.

#### Major fractions of Na_V_T, Na_V_P, and Na_V_R currents flow through Na_V_1.6 channelss

In neurons with resurgent currents, such as cerebellar Purkinje cells, Na_V_1.6 channels can carry all three current modes (Raman et al., 1997). Purkinje neurons in Na_V_1.6-null mice have reduced Na_V_T and Na_V_P current, and almost no Na_V_R current (Raman and Bean, 1997; Raman et al., 1997; Khaliq et al., 2003; Do and Bean, 2004). We tested for Na_V_1.6 contributions to the Na_V_T, Na_V_P, and Na_V_R current components using the Na_V_1.6 blocker, 4,9-ah-TTX, at a dose (100 nM) that is ∼10-fold higher than the IC50 (IC50 = 8 nM; Rosker et al., 2007) and still highly selective for Na_V_1.6 relative to other subunits.

In Na^+^-reduced external solution, 100 nM 4,9-ah-TTX blocked approximately 70% of I-Na_V_T in VGNs (Fig. 3A, B). Control and blocked (i.e., 4,9-ah-TTX sensitive currents) currents had similar V_1/2, Act_ and V_1/2, Inact_ values (Table 4). This is not surprising, given that the blocked current makes up most of the total current. V_1/2,Inact_ was, however, more negative for the unblocked (residual, 4,9-ah-TTX insensitive) current than for control current; V_1/2,Act_ was not significantly different (Table S1). This suggests the possibility of a second TTX-sensitive current that is not carried by Na_V_1.6 channels and has a more negative inactivation voltage dependence. The voltage dependencies of inactivation and activation of the two TTX-sensitive, transient conductances (blocked putative Na_V_1.6 and residual current) are consistent with observations on isolated rat VGNs (Liu et al., 2016).

**Figure 3.**
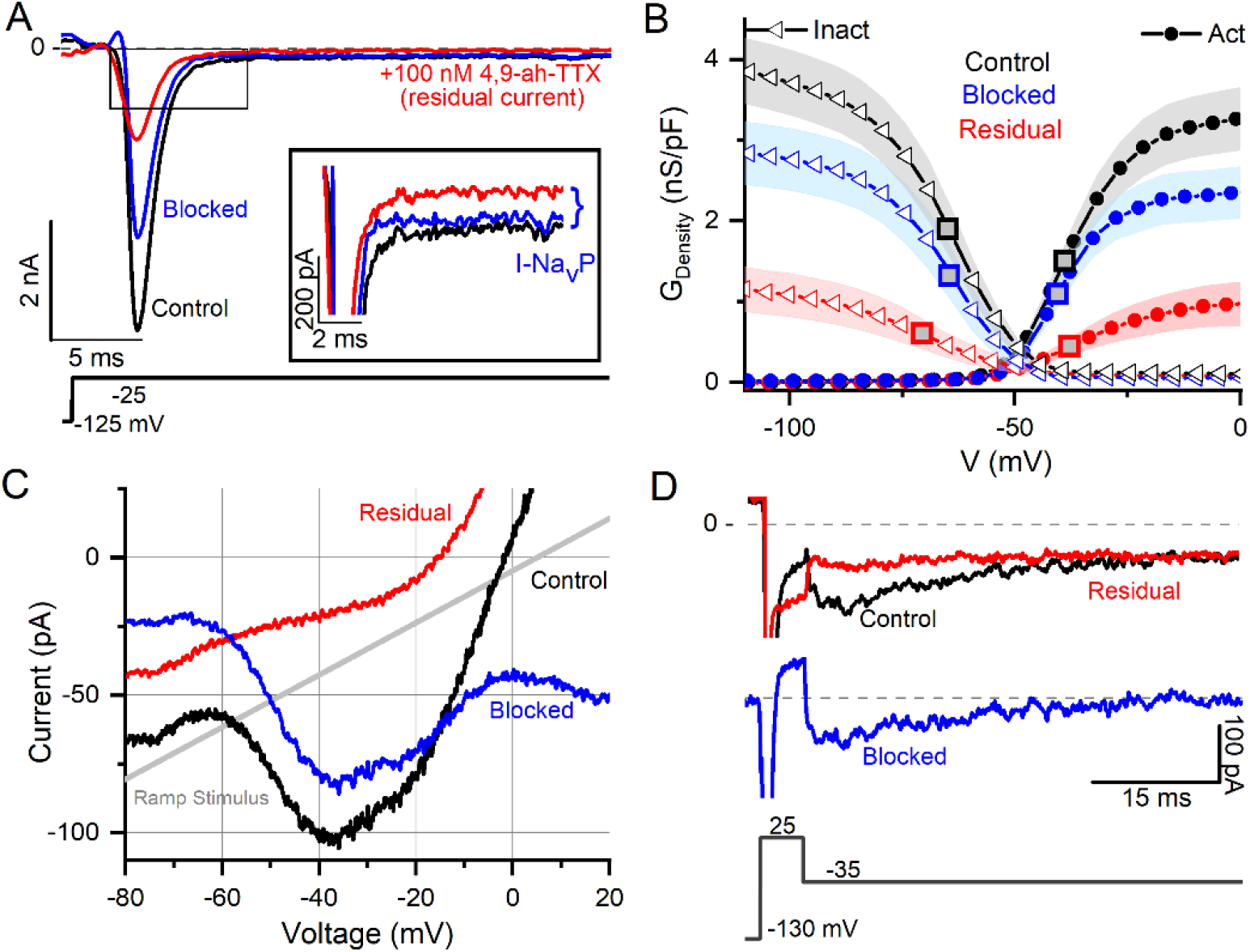
Na_V_1.6-selective channel blocker (4,9-ah-TTX) reveals strong Na_V_1.6 contribution to Na_V_T, Na_V_P, and Na_V_R current modes. **(A)** I-Na_V_T (P6 VGN) by 100 nM 4,9-ah-TTX is blocked by ∼70% (n = 13). Inset highlights block of Na_V_P current during voltage step. Control current is in black, red is the current measured during the drug (i.e., the “residual” current), and the blue is the blocked current (red current subtracted from the black). **(B)** Boltzmann fits of G-V activation and inactivation curves for data from (A). Square symbols indicate V_1/2_ values. 4,9-ah-TTX-insensitive current (red) had more negative inactivation V_1/2_ (n = 12, p = 0.02, Table 4). **(C & D)** Na_V_P (C, P17 VGN) and Na_V_R (D, P18 VGN) currents are also blocked (∼90%) by 100 nM 4,9-ah-TTX.

Na_V_P and Na_V_R currents were also blocked by 100 nM 4,9-ah-TTX. Block of I-Na_V_P was seen in responses to voltage steps (Fig. 3A inset) or to the slow voltage ramp (Fig. 3C) (n = 13). We were able to test 4,9-ah-TTX on just 1 of the 6 VGNs with Na_V_R; it produced a strong block of I-Na_V_R (Fig. 3D). These results with a Na_V_1.6-selective blocker suggest that Na_V_1.6 channels carry the majority of I-Na_V_ in cultured VGNs, including ∼50-70% I-Na_V_T and >90% of I-Na_V_P and most of I-Na_V_R. Similar results were obtained in calyx terminals (Meredith and Rennie, 2020).

#### I-Na_V_T and I-Na_V_P were enhanced by Na_V_ channel agonist ATX-II

The sea anemone toxin ATX-II interacts with Na_V_ channel gating, slowing down or preventing inactivation and thereby increasing Na_V_ current (Oliveira et al., 2004). ATX-II enhances Na_V_P current in vestibular afferent calyces (Meredith and Rennie, 2020) and Na_V_R and Na_V_P currents in spiral ganglion neurons (Browne et al., 2017) and dorsal root ganglion neurons (Klinger et al., 2012). We tested the impact of ATX-II on I-Na_V_ modes in mouse VGN cell bodies.

100 nM ATX-II increased maximum I-Na_V_T in 3 of 7 VGNs tested (Fig. 4A). In all VGNs, inactivation of I-Na_V_T was slowed, resulting in increased I-Na_V_P at the end of depolarizing steps (Fig. 4A) and during voltage ramps (Fig. 4B). On average, I-Na_V_P increased more than 5-fold (Fig. 4C); the example in Figure 4B was the largest effect seen. We detected no significant difference with ATX-II in V_1/2, Act_ and V_1/2, Inact_ values for I-Na_V_P and I-Na_V_T (Table 4, Table S1). For I-Na_V_T, ATX-II increased the slope factor (decreased the steepness of voltage dependence) of inactivation (p = 0.009, Hedges’ g = 0.85, large effect) but not activation (p = 0.85, power = 0.05) (not shown). We were not able to test ATX-II on resurgent current due to its low incidence.

**Figure 4.**
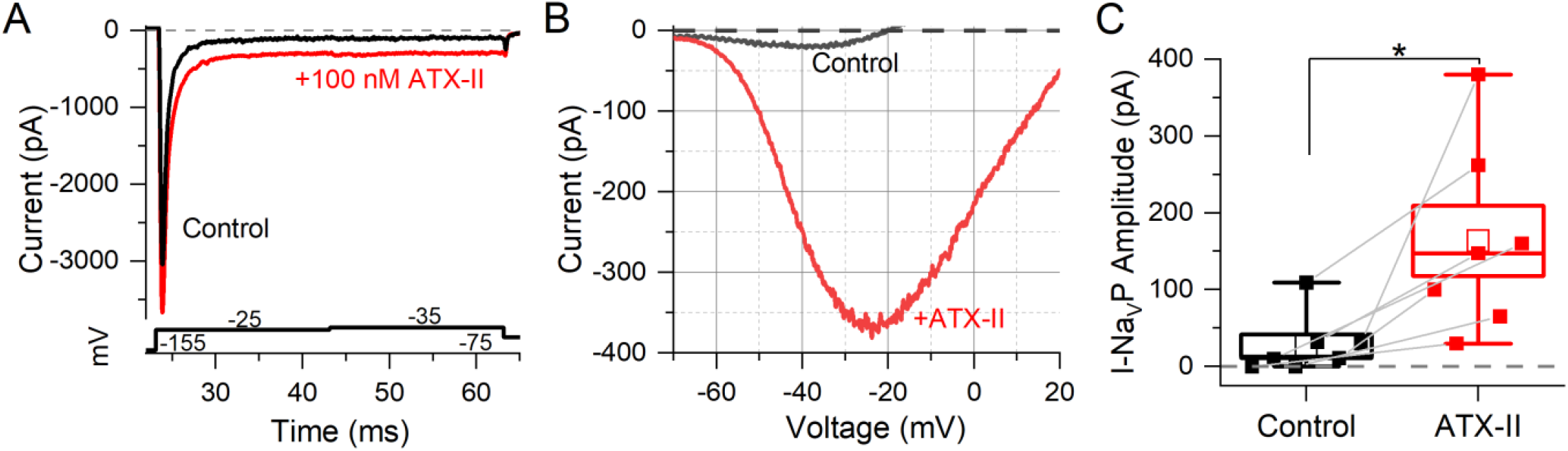
Na_V_ channel agonist (ATX-II) strongly enhanced Na_V_ T and Na_V_ P currents. **(A)** An example of enhanced Na_V_T and Na_V_P current by 100 nM ATX-II (red trace) (top panel) as elicited by depolarizing voltage steps (bottom panel). **(B)** I-V relation of Na_V_P current in a P7 VGN, before (grey) and in 100 nM ATX-II (red). Voltage dependence of I-Na_V_P was not significantly altered by ATX-II: V_1/2_ of G-V curves –38.8 ± 4.2 mV (n = 7) vs. –30.6 ± 2.9 mV, p = 0.12, power = 0.33 (not shown). **(C)** Na_V_P currents grew in ATX-II from 27.4 ± 14.4 pA to 163.5 ± 45.9 pA (n = 7, paired t-test; p = 0.01; Hedges’ g = 0.93, large effect). Na_V_R currents were not tested.

### 3.2 Roles of Na_V_ currents during action potentials and spike trains

To characterize AP waveforms and evoked firing patterns, we recorded in current clamp mode from 62 VGNs in K^+^ based standard solutions (Table 1). Without reduced external Na^+^, voltage clamp was less satisfactory: the large fast I-Na_V_ often escaped voltage clamp, obscuring the small, non-inactivating Na_V_P and Na_V_R currents. Although we could not identify I-Na_V_P or I-Na_V_R in whole-cell voltage clamp recordings, we characterized some features of I-Na_V_T and tested for effects of the Na_V_ channel blocker and agonist on spike waveform and firing pattern.

#### Na_V_ conductance correlated with features of the action potential waveform

We classified VGNs into four groups based on firing patterns evoked by depolarizing current steps just above current threshold (± 50 pA), following the scheme of Ventura and Kalluri (2019) (Fig. 5A). Transient neurons fired 1 or 2 spikes at step onset independent of step size. Sustained-firing neurons ranged from sustained-A type, with spike trains lasting the 500-ms depolarizing step, to sustained-B type, with shorter trains, to sustained-C type, with 2 or more small spikes that devolve into voltage oscillations. Accommodation could increase as more current was injected (Kalluri et al., 2010).

**Figure 5.**
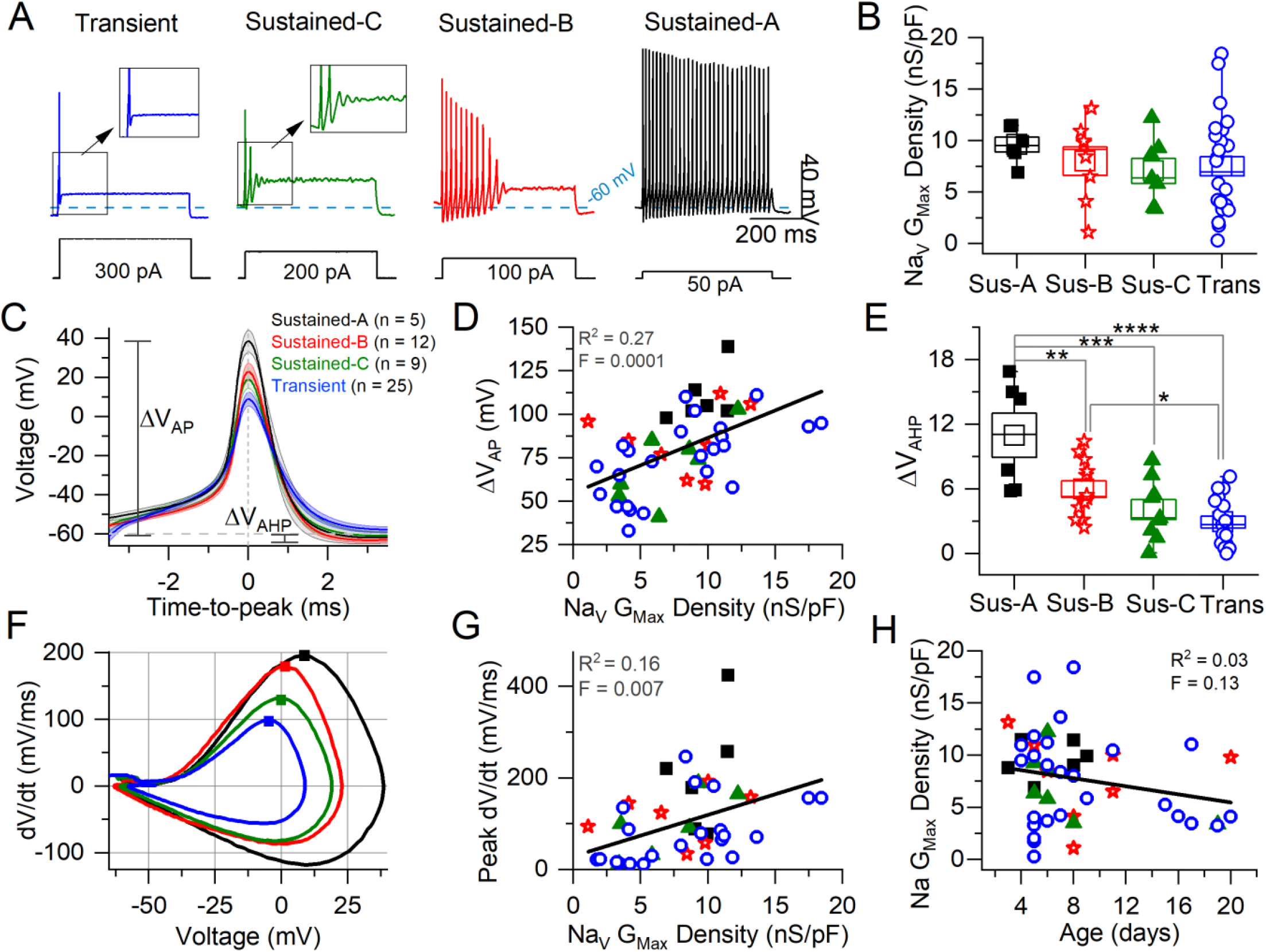
Correlating VGN firing patterns with spike waveforms and maximum Na_V_conductance density. **(A)** Exemplar firing patterns in VGNs evoked by a 500-ms current steps of different size reflecting different current thresholds. **(B)** Variation in maximum Na_V_ conductance density with firing pattern is not significant (homogeneous variance, Levene’s Test). **C:** Average spikes for each firing pattern, aligned to peak; spike height (ΔV_AP_) and afterhyperpolarization (AHP) are measured from V_rest_. **(D)** AP height correlated weakly with Na_V_ G_max_ density. **(E)** AHP depth varied with firing pattern (****, p = 0.00007; ***, p = 0.0005; **, p = 0.009; *, p = 0.04). **(F)** Phase plane plots of averaged APs from C. Squares denote peak dV/dt (rate of spike rise) values. **(G)** Peak dV/dt (see squares in F) correlated weakly with Na_V_ G_Max_ density. **(H)** No relationship between Na Gmax Density and age.

Transient VGNs had a significantly higher current threshold for spiking relative to sustained-A and sustained-B VGNs but we detected no significant difference in resting potential (V_rest_), input resistance (R_in_), or membrane capacitance (C_m_) across firing patterns (Table S2). While the incidence of the transient firing pattern was stable with age, at ∼50%, the distribution of sustained firing patterns changed from an approximate balance across sustained-A, -B, and -C subtypes for ages below P10 (n = 21) to mostly sustained-B above P10 (6/7) (Fig. S3). Similar changes with age were reported in rat VGNs (Ventura and Kalluri, 2019).

We assessed the AP waveform associated with each firing pattern (Fig. 5C, summarized in Table 5, and detailed in Table S3). APs from transient VGNs were smaller than APs from sustained-A VGNs (ΔV_AP_, Fig. 5C) and had slower peak rates of depolarization (peak dV/dt) than APs from sustained-A and sustained-B VGNs (Fig. 5F). Time-to-peak and voltage threshold of APs did not differ significantly across firing patterns (Fig. 5C, D).

**Table 5.** AP waveform differences between firing patterns from Figure 5. (one-way 4-factor ANOVA)

| Table 5: AP waveform difference between firing patterns (one-way 4-factor ANOVA) |  |  |  |  |  |  |
| --- | --- | --- | --- | --- | --- | --- |
| Firing pattern<br>(n) | Time-to-peak<br>(ms) | AHP<br>(mV) | $V_{AP}$<br>(mV) | Voltage<br>threshold (mV) | Peak dV/dt<br>(mV/ms) | $Na_V$ $G_{max}$ Den<br>(nS/pF) |
| Sustained-A<br>(6) | 4.5 ± 0.4 | 11.0 ± 2 | 110 ± 6.2 | -41.0 ± 3.3 | 208.3 ± 52.2 | 9.6 ± 0.7 |
| Sustained-B<br>(13) | 4.7 ± 0.3 | 6.0 ± 0.8 | 83.5 ± 6.4 | -44.3 ± 1.1 | 166.3 ± 34.2 | 8.0 ± 1.4 |
| Sustained-C<br>(10) | 4.5 ± 0.3 | 4.1 ± 0.9 | 80.0 ± 7.7 | -44.0 ± 1.3 | 123.7 ± 28.3 | 7.0 ± 1.2 |
| Transient<br>(28) | 4.2 ± 0.2 | 2.9 ± 0.6 | 75.1 ± 4.0 | -42.9 ± 1.3 | 78.6 ± 12.5 | 7.5 ± 0.9 |
| p | 0.45 | 0.00001 | 0.009 | 0.16 | 0.005 | 0.68 |
| power | 0.23 | 0.99 | 0.83 | 0.16 | 0.88 | 0.14 |

For 56 cells, we also collected Na_V_ currents in voltage clamp and fit activation (peak G-V) curves (Eq. 1) to measure maximum Na_V_ conductance density (Na_V_ G_Max_ density) (Fig. 5B). We had hypothesized that Na_V_ G_Max_ density would be highest for sustained-A VGNs and lowest for transient VGNs but actually did not detect a significant difference in Na_V_ G_Max_ density across firing pattern populations, although this was a low-powered observation (Fig. 5B, Table 5). Spike height (Fig. 5C) and peak dV/dt from phase plane plots (Fig. 5F; Bean, 2007) both correlated with Na_V_ G_Max_ density (Fig. 5D, G), as expected given that Na_V_ currents drive the rising phase of the AP. We saw no relationship between Na_V_ G_Max_ density and age (Fig. 5H).

The spike afterhyperpolarization (AHP) was significantly greater in sustained-A neurons than any other firing groups (Fig. 5E, Table 5, Table S3). The AHP preserves sodium channel availability by relieving inactivation, shortening the refractory period, and allowing sustained and regular firing at high rates (Gittis et al., 2010). Differences in AHP have previously been attributed to differences in K_LV_ conductance that affect resting potential, current threshold, and membrane recovery time (Hight and Kalluri, 2016; Kalluri et al., 2010; Iwasaki et al., 2008). In particular, the low K_LV_ conductance of immature sustained-A VGNs can contribute by making V_rest_ less negative.

In summary, maximum Na_V_ conductance density did not clearly vary with firing pattern but correlated with features of the spike waveform. We had hypothesized that sustained-A VGNs would have the highest Na_V_ G_Max_ densities and transient VGNs the lowest densities but did not detect a significant difference in average Na_V_ G_Max_ density across firing pattern populations, except that the range of G_Max_ densities was much higher for the transient VGNs. Key features of the AP waveform – spike amplitude, AHP, and peak rate of depolarization – did positively correlate with Na_V_ G_Max_ density (Fig. 5).

#### Blocking Na_V_1.6 currents reduced excitability and altered AP waveform

We probed the effects of the Na_V_1.6 blocker 4,9-ah-TTX on spiking (Fig. 6), which greatly reduced Na_V_T, Na_V_P, and Na_V_R currents recorded in Na^+^ Reduced/Cs^+^ solutions (Fig. 3). In standard recording conditions, 100 nM 4,9-ah-TTX showed a similar percent block of I-Na_V_T for transient VGNs (50.1 ± 11.3%) and sustained VGNs (52.0 ± 5.2%). Given that Na_V_1.6 current expressed in HEK cells has similar 4,9-ah-TTX sensitivity (Denomme et al., 2020), Na_V_1.6 channels may carry most of the Na_V_ current in our isolated and cultured VGNs.

**Figure 6.**
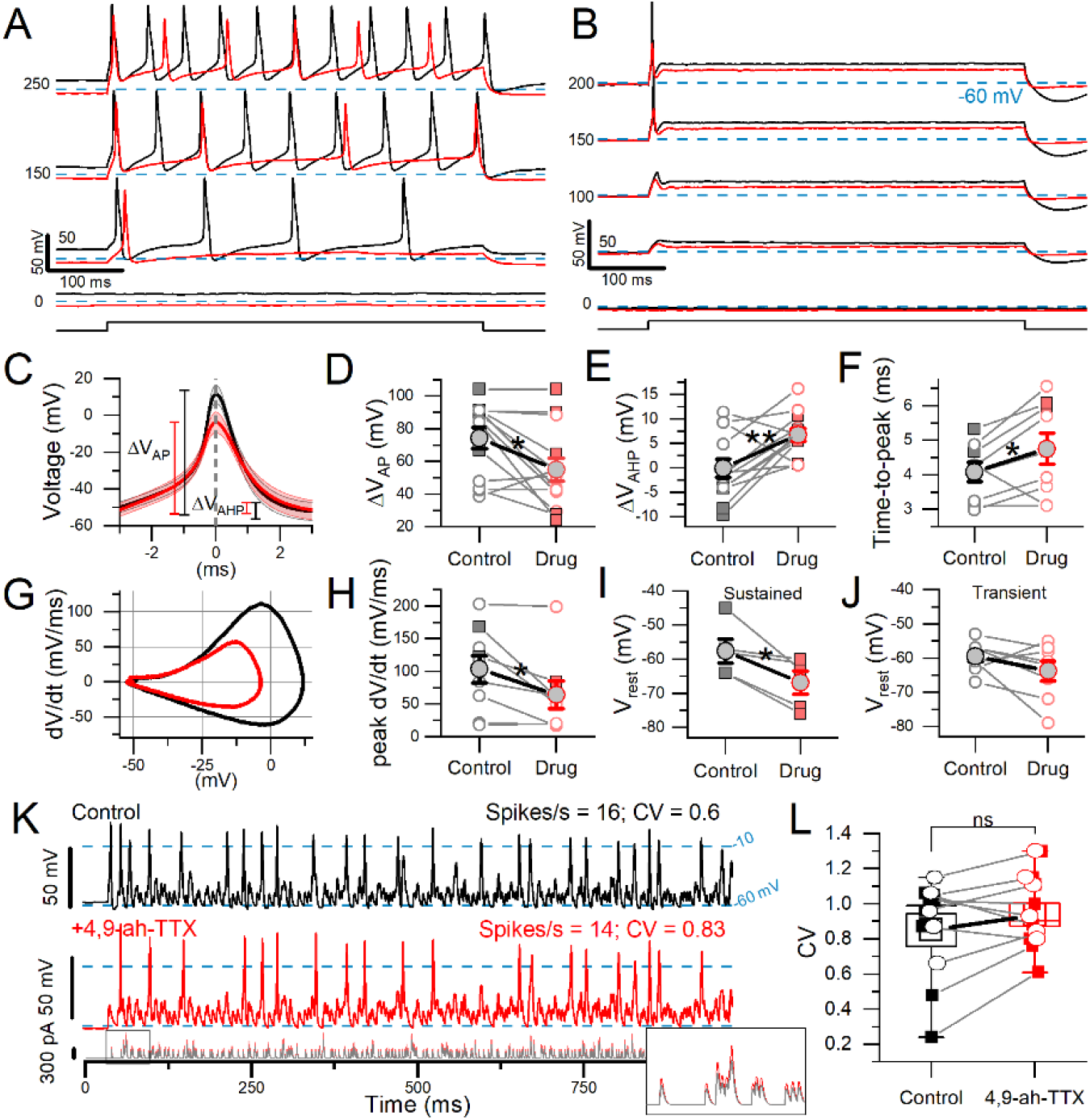
4,9-ah-TTX reduced step-evoked spiking excitability and altered the AP waveform. In 100 nM 4,9-ah-TTX, the 50% block of I-Na_V_T was similar in transient VGNs (n = 8) and sustained VGNs (n = 5: 1 sustained-A, 1 sustained-B, 3 sustained-C) (Welch’s t-test, p = 0.94, power = 0.05). **(A & B)** In 100 nm 4,9-ah-TTX (red), current threshold increased in a sustained VGN (A) and a transient VGN (B). In 7 cells, threshold increased from 92.3 ± 14.8 pA to 165.4 ± 23.6 pA (paired t-test, p = 0.002, Hedges’ g = 0.63; medium effect). **(C)** First AP from firing patterns such as (A) and (B), averaged (n = 7). **(D)** In 4,9-ah-TTX, for sustained (squares) and transient (circles) neurons, spike height decreased (p = 0.01), **(E)** afterhyperpolarization depth increased (p = 0.006), **(F)** and spike latency (i.e., time-to-peak) increased (p = 0.05). In some cases, AHP was depolarized relative to (V_rest_) by the current step. **(G)** Phase plane plots of mean AP waveforms in (C) highlight the change in peak dV/dt and V_AP_. **(H)** Peak dV/dt, from (E), was decreased by 4,9-ah-TTX (p = 0.03, Hedges’ g = 0.14, small effect). **(I & J)** Sustained VGNs (I) were significantly hyperpolarized (p = 0.03, Hedges’ g = 0.67, medium-large effect) but not transient VGNs (J) (p = 0.16). **(K)** 100 nm 4,9-ah-TTX (*red middle panel*) did not significantly reduce CV in spike trains evoked by pseudo-EPSCs (bottom panel) relative to control conditions (*black top panel*) in a P5 transient VGN. Dashed lines at –10 mV indicate event threshold for spike count; dashed lines at –60 mV indicate V_rest_. *Inset:* first 50 ms of EPSC train. **(L)** CV was not significantly altered in 4,9-ah-TX (paired t-test, 0.85 ± 0.09 vs 0.94 ± 0.07, p = 0.15, power = 0.29). Sustained VGNs (n = 4, *squares*); transient VGNs (n = 6, *circles*).

100 nM 4,9-ah-TTX affected step-evoked firing quantitatively, as shown in Figure 6. For all 13 VGNs tested, the Na_V_1.6 blocker increased current threshold for 500-ms steps. For sustained VGNs, 4,9-ah-TTX reduced the number of APs (spike rate) at spiking threshold and throughout the family of current steps (Fig. 6A). For transient VGNs, 100 nM 4,9-ah-TTX increased current threshold for spiking and decreased spike amplitude (Fig 6B).

To assess changes in the AP waveform, we temporally aligned the peaks of the first APs evoked by long current steps (Fig. 6C, detailed in Table S4). 4,9-ah-TTX reduced spike height by ∼20 mV (∼25%) on average (Fig. 6D). AHP depth was also reduced, possibly because hyperpolarizing K^+^ currents were less activated during the smaller spike (Fig. 6E) and AP time-to-peak increased (Fig. 6F). No change was detected in spike width at half-height, although control spikes are narrower at their peaks, reflecting their higher rates of depolarization and repolarization (Fig. 6G, H).

4,9-ah-TTX substantially hyperpolarized V_rest_ in sustained VGNs (Fig. 6I) but not transient VGNs (Fig. 6J). These data suggest that in sustained VGNs, 4,9-ah-TTX blocks depolarizing channels that are open at rest, such as I-Na_V_T and I-Na_V_P. The larger impact of the blocker on sustained VGNs may reflect a different balance of channels open at rest: they have smaller K_LV_ conductances (Kalluri et al., 2010) and may also have larger Na_V_ conductances open at rest, leading to a relatively more depolarized V_rest_.

To test the impact of Na_V_1.6 current on spike regularity (CV), we stimulated firing with frozen trains of synthetic (“pseudo”) EPSCs. The pseudo-EPSCs had pseudo-random timing to represent noisy quantal input from hair cells to afferent terminals where spiking normally initiates (Kalluri et al., 2010). In 4 of 14 VGNs tested, block of Na_V_1.6 current with 4,9-ah-TTX eliminated EPSC-induced spiking entirely. In the remaining 10 VGNs (6 transient and 4 sustained), we measured regularity as coefficient of variation (CV). To avoid rate effects on CV, we held spike rates to control levels (15.1 ± 1.3 vs 15.3 ± 1.7 spikes/s) by controlling EPSC size (Fig. 6K; see Kalluri et al., 2010). However, blocking Na_V_1.6 current did not significantly affect CV (Fig. 6L).

In summary, in all firing pattern groups, blocking Na_V_1.6 current with 4,9-ah-TTX altered spike wave form, decreasing spike amplitude, AHP, time-to-peak, and rate of depolarization. In sustained VGNs alone, 4,9-ah-TTX also made resting potential appreciably more negative, showing that Na_V_1.6 conductance is a significant factor in setting resting potential. Later (see Fig. S4), we use the computational model to assess the effects of individual current modes on V_rest_. In a sample of transient and sustained neurons, blocking Na_V_1.6 current had no consistent effect on spike regularity when overall rate was held constant by increasing EPSC size.

#### ATX-II increased excitability and spiking regularity

In voltage clamp, 100 nM ATX-II, which reduces Na_V_ channel inactivation, increased Na_V_P and Na_V_T currents (Fig. 4). We tested its impact on step-evoked firing patterns and pseudo-EPSC-evoked spike trains (Fig. 7).

**Figure 7.**
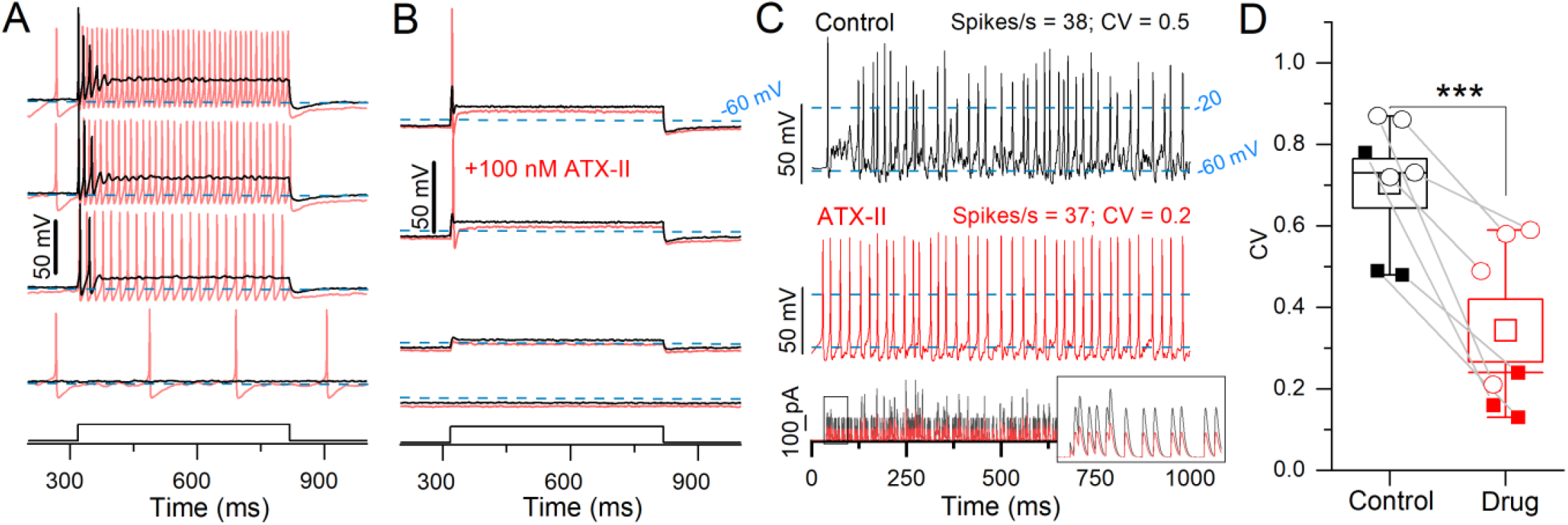
ATX-II increased spike rate and rate-independent regularity for current steps and EPSC-train stimuli. **(A)** 100 nm ATX-II (*red*) increased excitability relative to control (*black*) in a P12 sustained-C VGN: increasing the number of APs per step and inducing spontaneous spiking at rest (step current threshold = 0). **(B)** In a P12 transient VGN, current threshold was reduced, but the number of APs remained unchanged. (**C**) Pseudo-EPSC trains (*bottom pane*l) evoked spikes in the sustained-C VGN from A. ATX-II increased spike timing regularity in all VGNs. Inset shows the first 50 ms of EPSC train. To keep spike rate constant between conditions at ∼38 spikes/s, smaller EPSCs were applied during ATX-II (*bottom panel*). **(D)** ATX-II decreased CV in every VGN tested (transient = open circles, sustained = filled squares) (0.70 ± 0.06 to 0.34 ± 0.08, paired t-test; n = 7; p = 0.003).

In a sample of 9 VGNs, ATX-II reduced current threshold to zero and changed step-evoked firing patterns toward more sustained categories in 7 VGNs (3 sustained-A, 2 sustained-C, 2 transient); e.g., sustained-C VGNs became sustained-A (Fig. 7A). ATX-II also increased spike rate and decreased CV in responses to trains of pseudo-EPSCs. To control for rate effects on CV, we decreased EPSC amplitude in ATX-II to match spike rate (35 ± 3.4 sp/s) to the control value (31 ± 2.8, Fig. 7C) and still found a consistent, if modest, decrease in CV in ATX-II (Fig. 7C, D). Figure 7B shows one of 2 of 9 VGNs, both transient, that remained transient, irregular, and relatively inexcitable in ATX-II. There was no significant effect on AP waveform (Table S4).

In summary, reducing Na_V_ channel inactivation with ATX-II lowered current thresholds for both step-and pseudo-EPSC-evoked firing, in most cases to zero current (spontaneous firing). ATX-II also slightly increased spike regularity independent of rate. These results are consistent with increased Na_V_ channel availability near resting potential, presumably through reduced proportion inactivated channels, effectively increasing the persistent current.

### 3.3 Modeling the effects of transient, persistent, and resurgent Na_V_ currents on VGN firing

We hypothesized that I-Na_V_P and I-Na_V_R increase excitability in VGNs by increasing channel availability near spike threshold, and thus enhance the likelihood of firing. Because excitability has been associated with regularity in VGNs, we further hypothesized that I-Na_V_P and I-Na_V_R enhance spike timing regularity. Lacking pharmacological tools to disentangle the impacts of Na_V_T, Na_V_P and Na_V_R currents *in vitro*, we adapted existing models of neuronal firing to create model VGNs (“mVGNs”) for which each current mode could be adjusted. Below we show that, while I-Na_V_T affected rate and regularity in all mVGNs, I-Na_V_P significantly increased rate and rate-independent regularity in the sustained mVGNs and I-Na_V_R increased rate but decreased rate-independent regularity in the transient mVGNs.

We combined a single-compartment conductance-based VGN spiking model (Hight and Kalluri, 2016; Ventura and Kalluri, 2019) with equations for I-Na_V_P and I-Na_V_R (Wu et al., 2005; Venupogal et al., 2019). Our recorded range of Na_V_ conductance density (g_NaVT_) values was 0.2 – 18.4 nS/pF or 0.3 – 20.5 mS/cm^2^. We refined the spiking parameters for each mVGN by fitting the phase plane plot (PPP) of each mVGN to the PPP of the average recorded AP (Fig. 5F, see Methods). Based on the fitting results, we used 16 mS/cm^2^ for sustained-A, 13 mS/cm^2^ for sustained-B, 11 mS/cm^2^ for sustained-C and, and 7 mS/cm^2^ for transient model neurons (Table 3). These values neatly replicated AP waveforms, firing patterns, spike rates, and regularity ranges of each VGN type.

To test the impact of Na_V_ current components, we separately simulated four combinations of Na_V_T, Na_V_P and Na_V_R current modes of I-Na_V_ to assess differential effects on firing: 1) “T”, 2) “T + P”, 3) “T + R”, and 4) “T + P + R”. I-Na_V_P and I-Na_V_R were simulated with conductance density values as a percentage of I-Na_V_T; I-Na_V_P at 3% I-Na_V_T and for I-Na_V_R at 10% I-Na_V_T. These proportions are based on our mouse VGN data and values from calyx afferent terminal recordings in semi-intact vestibular organs (Meredith & Rennie, 2020) (Methods and Table 3). Current-voltage relations of the simulated currents reproduce the voltage dependence of experimental data (dashed curves in Fig. 1D).

Another set of simulations explored whether any changes with adding I-Na_V_P and I-Na_V_R were simply the effect of increasing Na_V_T conductance (g_NaVT_). We increased g_NaVT_ (“T+”) to match total conductances achieved in other simulations by adding Na_V_P and/or Na_V_R conductances, and to explore the range of g_NaVT_ recorded (Fig. 5B).

#### Adding I-Na_V_R or I-Na_V_P increased instantaneous firing rates and altered spike waveforms evoked by current steps

We tested responses of mVGNs to 500-ms current steps for comparison with step-evoked spike patterns (illustrated for real VGNs in Figure 5A). In Figure 8A, we show, for the 5 I-Na_V_ combinations, the first 50 ms of the response to a current injected after a 500-ms holding period (0 injected current). Individual Na_V_ current modes during spiking are shown in Figure 8B – D. To highlight any effects on spike waveform, Figure 8 (E, F) shows the waveforms and phase-plane plots of aligned first spikes of each mVGN firing pattern shown above. These simulations show that adding I-Na_V_R and I-Na_V_P increased excitability in sustained mVGNs.

**Figure 8.**
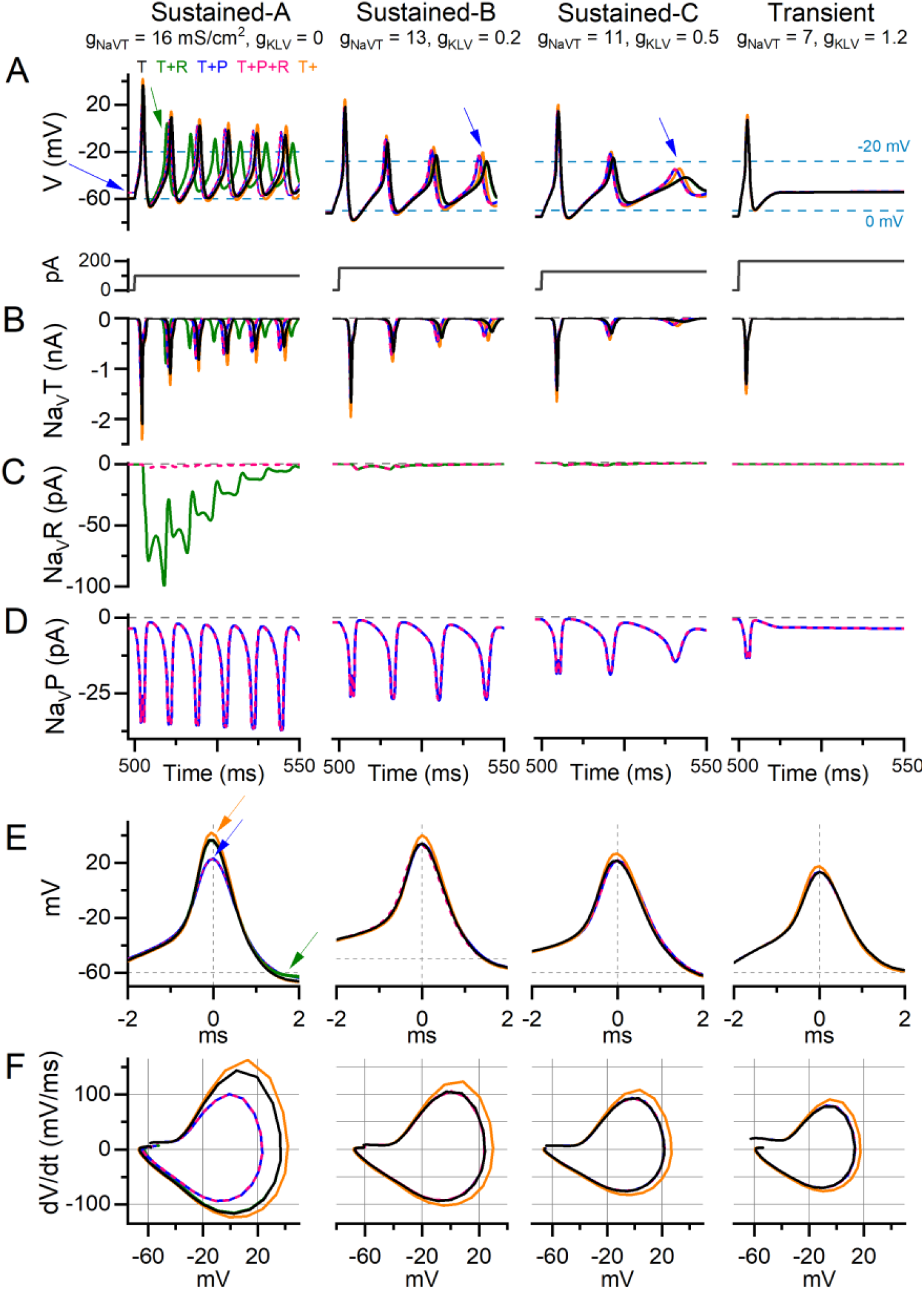
Adding I-Na_V_P altered AP waveform and ISI in current-step responses of model VGNs. Step-evoked firing in model VGNs with only I-Na_V_T (T); added I-Na_V_R (T+R) (10% of I-Na_V_T); added I-Na_V_P (T+P) (3% of I-Na_V_T); both added (T+P+R); and increased I-Na_V_T (T+). 1st 50 ms of responses to current step. Current steps, below, were increased (left to right) to account for increased current thresholds for spiking as firing patterns progressed from sustained-A to transient. Rows B-D show currents corresponding to the above firing pattern and isolated for each current mode (T, R, P). **(A)** In sustained-A, -B, and –C models, adding I-Na_V_P had variable effects on refractory periods and therefore AP time-to-peak in sustained-A, -B, -C mVGNs, and increased the size of post-spike voltage oscillations in sustained-B and sustained-C mVGNs (blue arrows). Adding I-Na_V_P depolarized Vrest in sustained-A mVGN (blue arrow), and slightly decreased refractory periods (time-to-peak) in all sustained mVGNs. **(B)** I-Na_V_T current flow during AP train. To control for increased Na_V_ conductance with added P and/or R conductance, we also ran a simulation with Na_V_T conductance density increased by the same amount (18.1, 14.7, 12.4, 7.9 mS/cm^2^ in sustained-A, -B, -C, transient simulations, respectively; T+ (orange) traces). **(C)** I-Na_V_R current flow during APs; note that I-Na_V_R flows during repolarization of each AP and decreases in amplitude with each successive AP in sustained-A model neuron. **(D)** I-Na_V_P current flow during AP train; note small variations in peak I-Na_V_P and the increase in amplitude with each AP in sustained mVGNs. **(E)** Temporally aligning the first APs from trains in (A) shows I-Na_V_P decreased spike height and I-Na_V_R decreases AHP (green arrows) in sustained-A, and increased Na_V_T increases spike height (orange arrows) in all mVGNs. **(F)** Phase plane plots show that increasing Na_V_T also increases peak dV/dt in all mVGNs and I-Na_V_P decreases peak dV/dt in sustained-S mVGN.

Adding I-Na_V_P produced a small inward current at 0 injected current, before spiking onset (Fig. 8D), in all mVGNs. This was largest in the sustained-A mVGN (Fig. 8A, left), where adding I-Na_V_P depolarized V_rest_ (blue arrow), reduced spike height (Fig. 8E, blue arrow) and rate of rise (Fig. 8F), thus altering the AP waveform (Fig. 8F). In both T+P and T+P+R conditions, the added I-Na_V_P affected the first spike in the train by decreasing time-to-peak by 11%, spike height by 19%, spike width by 20% and peak dV/dt by 30% (Table S5); adding I-Na_V_R (i.e., T+P+R) had no additional effect. I-Na_V_P also increased AHP by 36%, reducing I-Na_V_T inactivation and so leading to larger Na_V_T and P currents and driving larger spikes later in the train. I-Na_V_P had much smaller effects on AP waveforms of other mVGNs. In sustained-B and -C mVGNs, I-Na_V_P enhanced the number/rate and size of spikes and oscillations after the first spike (1 – 4% decrease in ISI; blue arrows, middle columns, Fig. 8A).

Effects of I-Na_V_R on spike waveform were first evident after a spike had occurred, as expected from Na_V_R current’s requirement for a large activating depolarization (AP upstroke) to block the channels, followed by relief of block by the AP downstroke (Fig. 8C; also Fig. 1D). In sustained-A mVGNs, adding I-Na_V_R to I-Na_V_T (i.e., T+R) decreased ISI, shortening AHP duration by 52% and reducing ISI by 16%, therefore changing instantaneous spike rate and changing the timing of individual spikes in the train (green arrow; Fig. 8A). I-Na_V_R had smaller or negligible effects in sustained-B, -C and transient mVGNs, likely because their smaller APs fail to reach the voltage range of “blocking” for I-Na_V_R (> +25 mV; Raman and Bean, 2001). This may also account for the dominance of I-Na_V_P current, such that T+P+R simulations resembled T+P simulations: depolarization of V_rest_ by I-Na_V_P yielded smaller APs, effectively eliminating the impact of I-Na_V_R (Fig. 8C, pink traces).

We compared the effects of adding Na_V_R and Na_V_P currents to the effect of increasing the Na_V_T conductance alone (i.e., T+; Fig. 8, orange traces). The effects on step-evoked firing of sustained mVGNs were not the same. Increasing I-Na_V_T decreased ISI after the first spike (by <1 ms, Table S5) less than either I-Na_V_P or I-Na_V_R (∼3%) (Fig. 8A, B). Increasing I-Na_V_T affected first-spike waveforms more than I-Na_V_R or I-Na_V_P, substantially increasing spike height and peak rate of rise of first-spike AP waveforms for all firing patterns (compare orange curves with all others, Fig. 8E, F).

In summary, adding I-Na_V_R and/or I-Na_V_P slightly increased instantaneous spike rate in step-evoked firing of sustained mVGNs. Increasing I-Na_V_T to the same total conductance affected spike height and rate of rise more than I-Na_V_R and/or I-Na_V_P.

#### Na_V_P current affected resting potential more than Na_V_T or Na_V_R currents

Adding I-Na_V_P depolarized the resting membrane potential in sustained-A mVGNs (Fig. 8A, blue arrow). To simulate the 4,9-ah-TTX block in real VGNs (Fig. 6), we simulated step-evoked firing in the sustained-A mVGN and transient mVGN with I-Na_V_T reduced by 70% and I-Na_V_P and I-Na_V_R by 90% (Fig. S4). Again, we set I-Na_V_P at 3% and I-Na_V_R at 10% I-Na_V_T. The simulated block replicated observations in real VGNs. Current thresholds increased and spike heights fell in both mVGNs. The diminution of the spike was most pronounced in the sustained-A mVGN, which also showed reduced spike rate and clearly hyperpolarized V_rest_. V_rest_ was negligibly affected in the transient mVGN.

We analyzed the influence of each Na_V_ current mode on the change in V_rest_ in sustained mVGN by varying P, R, and T combinations (Fig. S4C). Blocking I-Na_V_R had no effect on V_rest_ in any mVGN because resurgent current requires activity to activate. Blocking I-Na_V_T had at most a small effect on V_rest_ and then only in sustained mVGN (∼1 mV). Virtually all the effect on V_rest_ was through I-Na_V_P: 6 mV for sustained mVGN and 0.5 mV for the transient mVGN.

Therefore, reduction in Na_V_T, Na_V_P, and Na_V_R currents in mVGNs reproduced key effects of a Na_V_1.6 channel blocker and show I-Na_V_P driving 5-10 mV depolarization of V_rest_ in sustained-A mVGNs. This simulation indicates that even if Na_V_P currents are equally present in sustained and transient VGNs (as we have modeled them), the effect of Na_V_P current is substantially stronger in sustained VGNs.

#### Adding I-Na_V_T, I-Na_V_P, and I-Na_V_R increased spike rate evoked by pseudo-EPSC trains and differentially affected regularity

To examine the roles of Na_V_ current modes in spike rate and regularity, we drove sustained-A and transient mVGNs with 1-second trains of simulated synaptic events (pseudo-EPSCs, first 500 ms shown in Fig. 9A, B), modeled after EPSCs recorded from vestibular afferent calyx events and quasi-randomly distributed in time (see Methods). We generated a set of 5 different pseudo-EPSC trains and applied the same set to each simulation condition shown in Figure 9. Therefore, the input noise is similar across each g_NaVT_ value in Figure 9. Each measurement in panels C-H is the mean and SEM for the 5 pseudo-EPSC trains at the specified g_NaVT_ and combinations of Na_V_R and Na_V_P conductances. Figure 9B (bottom trace) shows one of the pseudo-EPSC trains, with an inset highlighting a short onset segment at high gain. Again, we held Na_V_P and Na_V_R conductance at 3% and 10% of g_NaVT_, respectively. Larger % values of Na_V_P are explored in Supplemental Figure 5.

**Figure 9.**
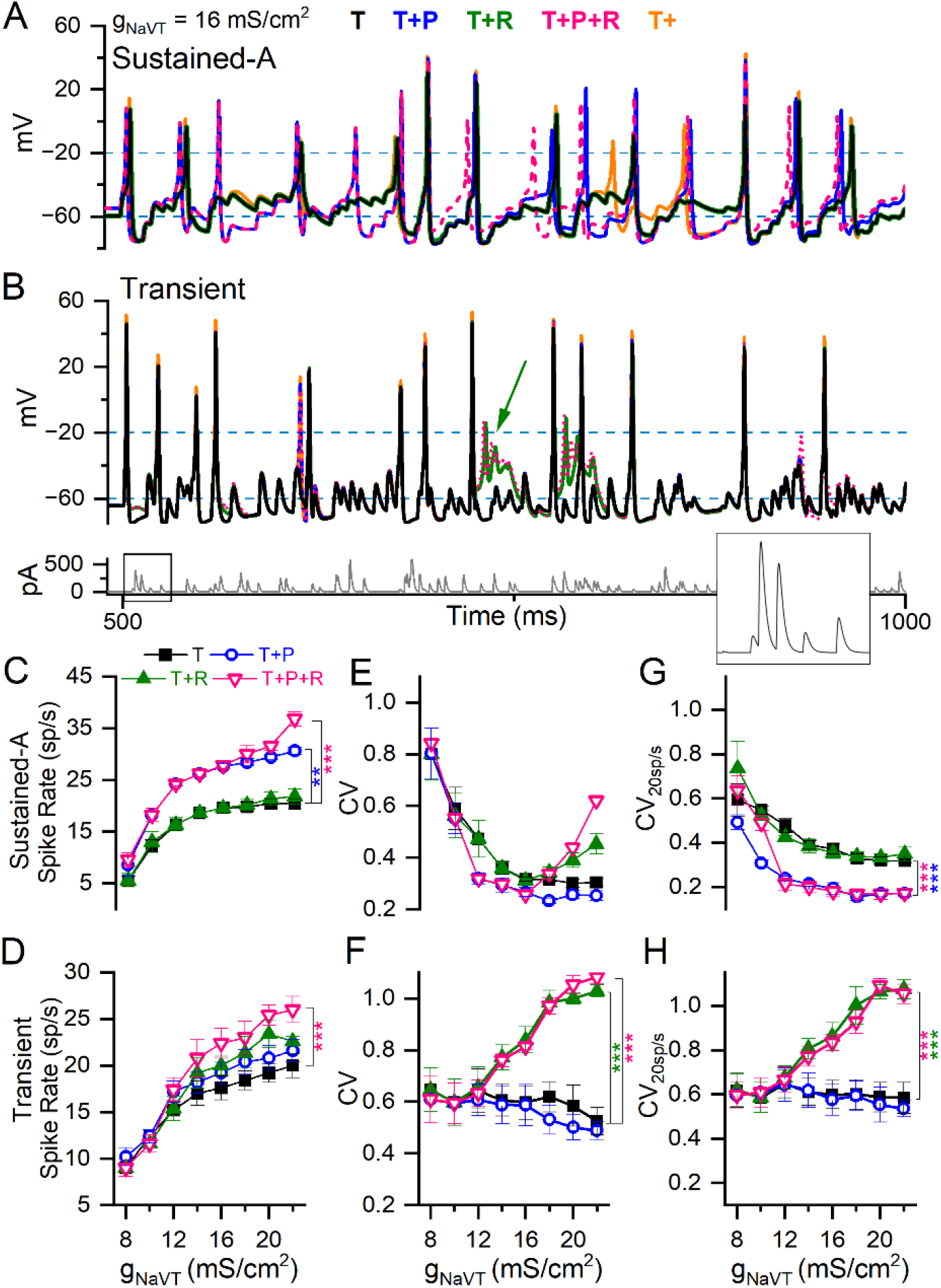
Adding I-Na_V_P and/or I-Na_V_R to I-Na_V_T differentially affected spike rate and regularity in model VGNs. **(A)** Sustained-A mVGN under different Na_V_ current combinations: adding I-Na_V_P increased spike rate, whether I-Na_V_R was also added or not. **(B)** Transient mVGN under different Na_V_ current combinations: adding I-Na_V_R increased spike rate, but sometimes caused depolarization block (green arrow). *Bottom panel*, pseudo-EPSC train used to evoke spike trains shown. **(C & D)** Summary of spike rates for each combination of Na_V_ current modes at each level of g_NaVT_. Increasing Na_V_T increased spike rate in both sustained-A and transient mVGNs. ***C***, Adding I-Na_V_P (T+P and T+P+R) increased spike rate relative to Na_V_T in sustained-A mVGN, while adding R has little effect on spike rate except at the highest value of (T+P+R). ***D***, I-Na_V_R (T+R and T+P+R) increased spike rate in transient VGN, while I-Na_V_P had no effect. **(E & F)** Spike train regularity (CV) for spike trains in C & D. I-Na_V_P had little effect on CV in either mVGN. I-Na_V_R increased CV especially in the transient mVGN, but also in sustained-A mVGN for g_NaVT_ > 14 mS/cm^2^. Two-way ANOVAs confirmed interactions between effects of g_NaVT_ and Na_V_ current modes on spike rate and CV in sustained-A and on CV in transient mVGNs (Spike rate: F_(8,24)_ = 3 and 0.8, p <0.0001 and 0.67. CV: F_(8,24)_ = 2 and 3.6 and p = 0.009 and <0.001). **(G & H)** CVs with spike rate held at ∼20 spikes/s (CV_20sp/s_) across conditions, used to assess the effects of Na_V_ currents on CV independent of spike rate. ***G***, Adding I-Na_V_P to sustained-A mVGN significantly decreased CV_20sp/s_ in sustained-A mVGN, with or without I-Na_V_R. ***H***, Adding I-Na_V_R to transient mVGN significantly increased CV_20sp/s_ in sustained-A mVGN for g_NaVT_ > 12 mS/cm^2^, with or without I-Na_V_P.

First, we assessed the effects of I-Na_V_T alone on spike rate and spike regularity at 12 values of g_NaVT_ density chosen to cover the range we encountered in VGN recordings (6 to 22 mS/cm^2^ in intervals of 2; orange traces in Fig. 9A and B). To study spike rate effects, we generated EPSC trains with average inter-event interval fixed at 5 ms (200 EPSCs/s) and an amplitude that produced ∼20 spikes/s at 14 mS/cm^2^, then varied g_NaVT_ density. This firing rate corresponds approximately to the measured background rate for mouse vestibular afferents (∼50 spikes/s, Lasker and Minor 2006) after compensation for the ∼15°C lower temperature of our room temperature VGN recordings. For all mVGNs, spike rate increased and CV decreased with increasing g_NaVT_ density (Fig. 9C – F), as previously demonstrated by Hight and Kalluri, 2016. For transient neurons, spike rate increased with g_NaVT_ density, again especially at the low end (Fig. 9D). But CV showed much less variation with g_NaVT_ density (Fig. 9F). Note that the sustained mVGN in the ‘T’ condition is no more regular than the transient mVGN for g_NaVT_ densities <12 mS/cm^2^ or more (compare panels E and F).

In a second set of simulations, we assessed CV at a constant spike rate of 20 spikes/s (CV_20sp/s_) to eliminate the confounding effect of spike rate on regularity. Rate was controlled by titrating the size of pseudo-EPSCs in each train, while not interfering with the pseudo-randomly distributed inter-event intervals. For the sustained-A mVGN (Fig. 9G), increasing g_NaVT_ density decreased CV_20sp/s_ up to about 12 mS/cms^2^, above which CV plateaued. For the transient mVGN (Fig. 9H), in contrast, there was no obvious change in CV over the entire range of g_NaVT_ tested. The addition of I-Na_V_R and/or I-Na_V_P to I-Na_V_T affected rate, CV and CV_20sp/s_ differently for the sustained-A mVGN vs. the transient mVGN (Fig. 9C-H).

#### Na_V_P and Na_V_R currents increase spike rate in sustained and transient mVGNs, respectively

For the sustained-A mVGN, rate increased substantially when Na_V_P current was added (p < 0.0001; Fig. 9C; Table S6) but was not affected by adding Na_V_R current except for the (T+P+R) condition at the highest level of g_NaVT_ density tested. For the transient mVGN, adding both I-Na_V_P and I-Na_V_R increased rate for g_NaVT_ density > 12 mS/cm^2^, reaching significance at the highest level. The effects of the two components appeared to be additive (Fig. 9D). In simulations of all three components (T+P+R), the increase in spike rate is compounded: sustained mVGNs showed a medium to large effect (Hedge’s g = 0.6) and transient mVGNs a small effect (Hedge’s g = 0.2) (Fig. 9C, D, Table S6).

In Supplemental Figure 5, we show the effect of increasing Na_V_P current to the highest values recorded at afferent terminals on hair cells, near the spike initiation zone: 5 to 10% of g_NaVT_ (Meredith and Rennie, 2020). Spike rate increased further as I-Na_V_P increased. For the sustained-A mVGN, spike rate doubled as I-Na_V_P increased from 0-10% (p < 0.0001, Hedges’ g = 1.5). For the transient mVGN, the impact on rate was smaller (∼25% increase; p < 0.001, Hedges’ g = 0.7).

#### I-Na_V_P decreases regularity in sustained mVGNs whereas I-Na_V_R increases regularity in transient mVGNs

Adding Na_V_P and Na_V_R conductances had complementary effects on the regularity (CV and CV_20sp/s_) of sustained-A and transient mVGNs. For the sustained-A mVGN (Fig. 9G), adding Na_V_P (at 3%) halved CV_20sp/s_ for all g_NaVT_ density > 8 mS/cm^2^, whether Na_V_R conductance was present or not. For the transient mVGN, in contrast, Na_V_P current had no detectable effect, but Na_V_R current increasingly enhanced irregularity (both CV and CV_20sp/s_) for g_NaVT_ density > 12 mS/cm^2^ (Fig. 9F, H). One factor in the enhanced irregularity is likely to be depolarization block at high total Na_V_ conductance (e.g., green arrow, Fig. 9B): instead of returning to V_rest_, the increased proportion of unblocked channels held V_m_ depolarized, leading to profound inactivation of Na_V_T current.

In summary, simulations suggest that I-Na_V_P and I-Na_V_R differentially affect spike rate and reinforce the difference in regularity at constant rate of sustained-A and transient VGNs. Adding P current made the sustained-A mVGN more regular, and adding R current made the transient mVGN more irregular, in both cases by factors of ∼2.

## 4 Discussion

### 4.1 VGN as a model preparation for firing patterns in the vestibular nerve

VGNs are cell bodies that, when mature and *in vivo*, are bipolar, inter-nodal and myelinated. Research from multiple groups (Chabbert et al., 1997; Limon et al., 2005; Iwasaki et al., 2008) has shown that isolated VGNs express voltage-gated currents and firing patterns that resemble currents and firing at spike initiation zones just below the hair cell-afferent terminal synapses in the vestibular epithelium, as illustrated by immunohistochemistry and direct recordings from calyx terminals (e.g., Lysakowski et al., 2011; Songer and Eatock, 2013; Meredith and Rennie, 2020). It is not clear whether the presence of these channels in the dissociated cell bodies represents true somatic expression, or whether nodal Na_V_ channels and paranodal K_V_ channels on either side of the soma come along during dissociation. In most cases, investigators have cultured the dissociated VGNs overnight, causing the VGNs to shed their myelin and expose the neuronal membrane for whole-cell patch recording.

In rat VGNs, there is some evidence that overnight culturing eliminates expression of some types of pore-forming Na_V_ channels. Dissociated young rat VGNs that were studied acutely had multiple transient Na_V_ currents with distinct voltage dependence and TTX sensitivity (Liu et al., 2016), including a Na_V_1.5 current (negatively shifted voltage range, TTX-insensitive) and a Na_V_1.8 current (positively shifted voltage range, TTX-resistant). These non-TTX-sensitive forms were not detected in overnight-cultured rat VGNs (Liu et al., 2016). Age differences across studies may also matter. Na_V_1.5 expression in ganglion cell bodies may be an immature feature, as suggested by its time course of expression in small rat DRG neurons (Ranganathan et al. 2002) and some decline in current expression over the first postnatal week in rat VGNs (Liu et al. 2016). Meredith and Rennie (2020) recorded TTX-insensitive currents (consistent with Na_V_1.5) from gerbil afferent calyceal terminals, and but only in afferents younger than P12.

In summary, isolated and cultured VGNs are compact cell bodies that allow high-quality voltage clamp recordings and reproduce some but not all naturally occurring ion channel expression. Importantly for our purposes, they show a range of firing patterns consistent with a role for intrinsic neuronal properties in setting up the different encoding strategies (temporal vs. rate) characteristic of irregular and regular afferent populations (Jamali et al., 2016; Cullen, 2019). Previous work has documented the difference in firing patterns of VGNs and probed their relationship to specific ion channels: low-voltage-activated K_V_1 (Iwasaki et al., 2008) and K_V_7 channels (Kalluri et al., 2010), Ca^2+^-dependent K channels (Limon et al. 2005), HCN channels (Horwitz et al., 2014) and Na_V_ channels (Liu et al. 2016). By going beyond correlation and developing a method to interrogate regularity in these synapse-less somata, a particularly strong case was made for K_LV_ channels (Kalluri et al., 2010). We took this approach with Na_V_ channel modes and provide support for a substantial role of relatively small persistent and resurgent currents in excitability and related properties such as resting potential.

### 4.2 Na_V_1.6 is the major Na_V_ subunit in VGNs, contributing to resting potential and excitability

All VGNs in our sample expressed I-Na_V_T and about half also expressed I-Na_V_P with no clear change in incidence with age (range P3-28), in contrast to mouse cochlear ganglion neurons where both persistent and resurgent forms increased with maturation (Browne et al., 2017). We saw no I-Na_V_R before P10. Beyond P10, the incidence was 12% (6/49 tested). VGNs also have voltage-dependent Ca^2+^ (Ca_V_) currents (L-, N-, P/Q-, R-, and T-type) (Desmadryl et al. 1997; Chambard et al. 1999) which drive Ca^2+^-dependent K^+^ currents that reduce sustained firing (Limón et al., 2005). Here we suppressed both Ca_V_ and Ca^2+^-dependent K^+^ currents by eliminating all but trace Ca^2+^ from the external medium. Our experiments do not indicate whether sustained VGNs have more Na_V_P or Na_V_R current than transient neurons, because the conditions for studying firing pattern did not allow isolation of the small P and R currents. However, our modeling suggests that for transient neurons, Na_V_P and Na_V_R currents, if present, would have little effect on regularity and resting potential.

RT-PCR of whole rat vestibular ganglia indicated expression of most Na_V_ pore-forming (α) subunits and all auxiliary (β) subunits (Liu et al. 2016). Block by the Na_V_1.6-selective blocker, 4,9-ah-TTX, however, indicates that Na_V_1.6 channels make up much or all of the current we recorded, for all modes, again consistent with results from calyx terminals in gerbil cristae (Meredith and Rennie, 2020). In our study, 100 nM 4,9-ah-TTX blocked Na_V_T current by a similar amount (∼50%) in sustained and transiently firing VGNs, suggesting that they express similar proportions of Na_V_1.6 channels. Persistent and resurgent I-Na_V_ were almost fully blocked. Cochlear spiral ganglion neurons also express 4,9-ah-TTX-sensitive transient, persistent, and resurgent currents (Browne et al. 2017), and are immunoreactive for Na_V_1.6 at spike initiation zones next to their terminals on hair cells (Kim and Rutherford, 2016). I-Na_V_R is theorized to arise from the internal block of Na_V_ channels, canonically Na_V_1.6, by a positively charged molecule such as Na_V_ auxiliary subunit β4 (present in RT-PCR data of the ganglia; Liu et al., 2016) or fibroblast growth factor homologous factor 14 (Raman and Lewis, 2014; White et al., 2019; Aman and Raman, 2024).

The residual Na_V_ current in 4,9-ah-TTX had significantly more negative inactivation than the control and blocked currents, but not nearly as negative as the TTX-insensitive current carried by Na_V_1.5 in acutely cultured, immature rat VGNs (Liu et al., 2016). RT-PCR data from rat vestibular ganglia (Liu et al., 2016) suggest Na_V_1.1, 1.2, 1.3, and 1.7 as candidates.

As expected for a major Na_V_ current, the Na_V_1.6 currents are critical to excitability in both firing types. In both sustained and transient neurons, spikes were substantially smaller in 4,9-ah-TTX, as expected from the strong block of the dominant transient mode of Na_V_1.6 current. A large decrease in AHP size presumably reflected both the small spike height (reducing activation of K^+^ currents that cause AHPs) and the more negative V_rest_ (reducing the driving force, i.e., voltage difference from E_K_). 4,9-ah-TTX blocking experiments showed that Na_V_1.6 channels contribute to resting potential in sustained neurons (Fig. 6I; Table S4).

Our modeling also suggests that Na_V_P and, to a lesser extent, Na_V_T current modes may significantly contribute to resting conductance in sustained VGNs (Fig. S4). Although I-Na_V_P contributed just 3% of maximum Na_V_ current density in VGNs, Na_V_P current at resting potential may be closer to 10% of total current, based on its relative voltage dependence and the substantial (∼30%) steady-state inactivation of Na_V_T current even at resting potential (Fig. 2B). The small to negligible effect of blocking Na_V_1.6 channels on resting potential in transient VGNs (Fig. 6J vs. 6I) is consistent with resting conductance being dominated by their greater density of K_LV_ channels. HCN conductances may also contribute to resting membrane potential in VGNs, depending on the balance of resting conductances (Ventura and Kalluri, 2019).

### 4.3 Effects of Na_V_ current modes on AP waveforms, firing patterns, rate, and regularity

AP height and rate of rise varied across firing pattern (Fig. 5C, F) and correlated modestly with maximum Na_V_ conductance density (Fig. 5D), both for all VGNs and for transient VGNs alone, which showed the greatest variance in maximum Na_V_ conductance density. Firing pattern, in contrast, did not correlate closely with Na_V_ G_max_ density. G_max_ density is dominated by the Na_V_T current mode, so these correlations do not reveal roles for I-Na_V_P, which has a more significant effect near resting potential, and I-Na_V_R, which acts during spike repolarization. To isolate their effects, we used simulations.

Our simulations support previous work on isolated VGNs indicating a dominant role for K_LV_ currents in differentiating spike rate and regularity of sustained (presumed regular) and transient (presumed irregular) VGNs (Kalluri et al., 2010; Ventura and Kalluri, 2019). K_LV_ currents increase irregularity independent of rate by enhancing the afferent’s sensitivity to high-frequency noise (Kalluri et al., 2010). We found that the large K_LV_ currents of transient VGNs limited the influence of Na_V_ current modes on rate and regularity, but when K_LV_ currents are small, as in sustained VGNs, augmenting transient and/or persistent Na_V_ currents increased excitability, spike rate, and rate-dependent regularity.

In sustained model VGNs, I-Na_V_P had a stronger effect than other modes on resting potential, depolarizing V_rest_ by ∼6 mV. The effect on V_rest_, in turn, shapes many AP metrics by affecting resting (input) conductance, inactivation state of I-Na_V_T, and current and voltage thresholds for spikes (Fig. 8F). I-Na_V_P also changed AP waveform by reducing time-to-peak and spike height in sustained-A mVGN APs. Primarily, however, I-Na_V_P strongly increased firing rate in sustained-A mVGNs. Regularly firing neurons, like sustained-A VGNs, use firing rate to preferentially encode low frequency information (Jamali et al., 2016; Cullen, 2019). A mechanism that increases firing at or near V_rest_ without inactivating will enhance ability to encode small excitatory and inhibitory inputs, increasing response linearity.

In sustained mVGNs current step simulations (Fig. 8A, B), adding I-Na_V_R altered instantaneous spike rate by truncating the AHP, decreasing the current threshold for spiking, and shortening ISI between the first two spikes. However, when tested with trains of pseudo-EPSCs, I-Na_V_R had no effect on average spike rate across multiple spike trains (Table S6), suggesting that long term effects of resurgent current reduce the impact of short-term changes in rate. In transient mVGNs, resurgent current affected spike rate and regularity (CV) during pseudo-EPSC trains, independent of spike rate and not. We attribute these effects to sensitization during the inter-spike interval of transient mVGNs to the pseudo-randomly times EPSCs.

The isolated VGN patch preparation and model provide an opportunity to isolate the impact of the afferents’ diverse voltage-gated channels, but incorporation into a fuller model of the inner ear may yield more insight into their impact on shaping the vestibular afferent signal. *In vivo*, many features differ across the peripheral and central epithelial zones that give rise to regular and irregular afferents, including the macromechanical input to hair cells, numbers and types of synaptic inputs, relative contributions of non-quantal and quantal transmission, size of dendritic arbors, and complements of expressed ion channels, all of which might shape voltage noise at spike initiation zones. Our analyses suggest that diverse Na_V_ currents flowing through one kind of pore-forming subunit can play significant roles in differentiating regular and irregular firing in the vestibular nerve.

## 6 Author Contributions

SBL conceived the project, planned, and performed the electrophysiological recordings and computational modeling, and wrote the first draft of the manuscript. RAE participated in planning, design, analysis, and writing the manuscript.

## 8 Acknowledgments

We are grateful to Dr. Omar López-Ramírez for sharing excitatory postsynaptic current data from vestibular afferent terminals and to Dr. Radha Kalluri for access to the original VGN spiking model. We thank Dr. Hannah Martin, Dr. Aravind Govindaraju, Dr. Kalluri, Dr. Daniel Bronson, Dr. Nathaniel Nowak, and Katherine Regalado for comments on early versions of the manuscript.

## 10 Data Availability Statement

The associated code is accessible in the following repository: https://github.com/eatocklab/NaV1025
currents-in-VGN-spiking.

The datasets generated and analyzed for both electrophysiology and modeling experiments can be
found in our Dryad repository:
https://datadryad.org/stash/share/sbukzLlDkLPlsuTAISqONfWVmLOuSwvnNNXr5LKdD6c
(Reviewer URL).

## 13 Supplementary Material

### 13.1 Supplementary Figures

**Figure S1.**
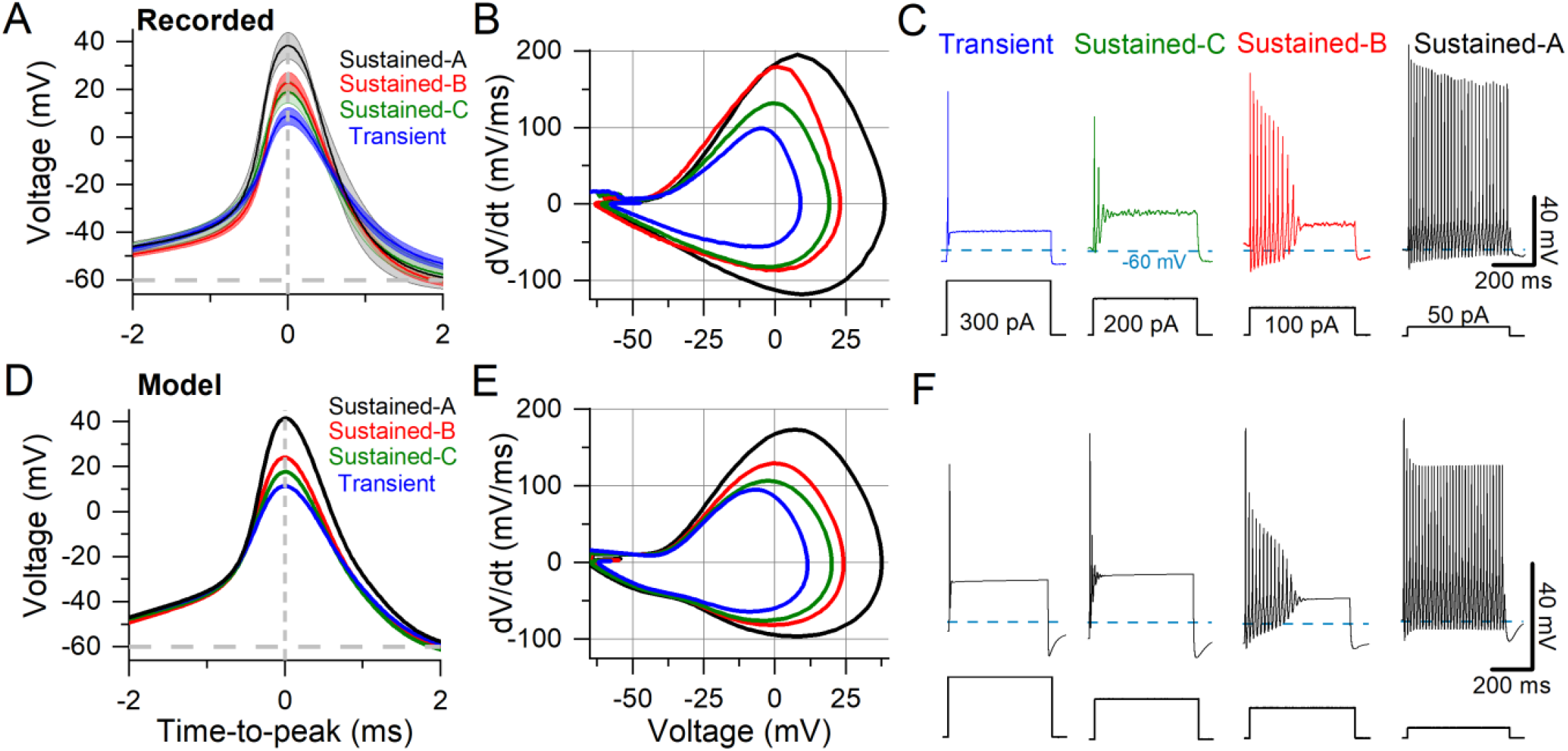
Comparison of VGN responses to model VGN simulations. **(A** – **C)** (reproduced from Fig. 5). Averaged recorded APs (A), corresponding mean phase plane plots (B), and exemplar step-evoked firing patterns (C). **(D)** Model-generated APs based on APs from (A). **(E)** Phase-plane plots for simulated APs from (D). Simulated APs have smaller peak rates of depolarization. **(F)** Step-evoked firing patterns of model VGNs capture key features of exemplar data in (C).

**Figure S2.**
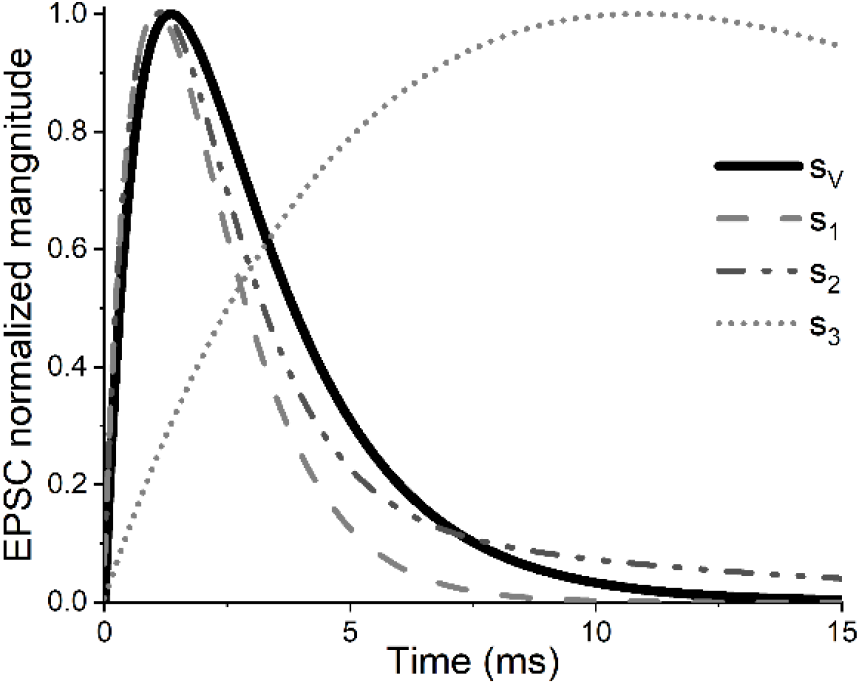
Shape of simulated EPSCs. Simulated excitatory postsynaptic currents (EPSCs) used to evoke spiking in our cell-autonomous model were modeled after spontaneous vestibular synaptic potentials (s_V_) recorded at room temperature in mouse calyces. EPSC shapes previously tested in Hight and Kalluri (2016) (s1, s2, and s3) are shown for comparison.

**Figure S3.**
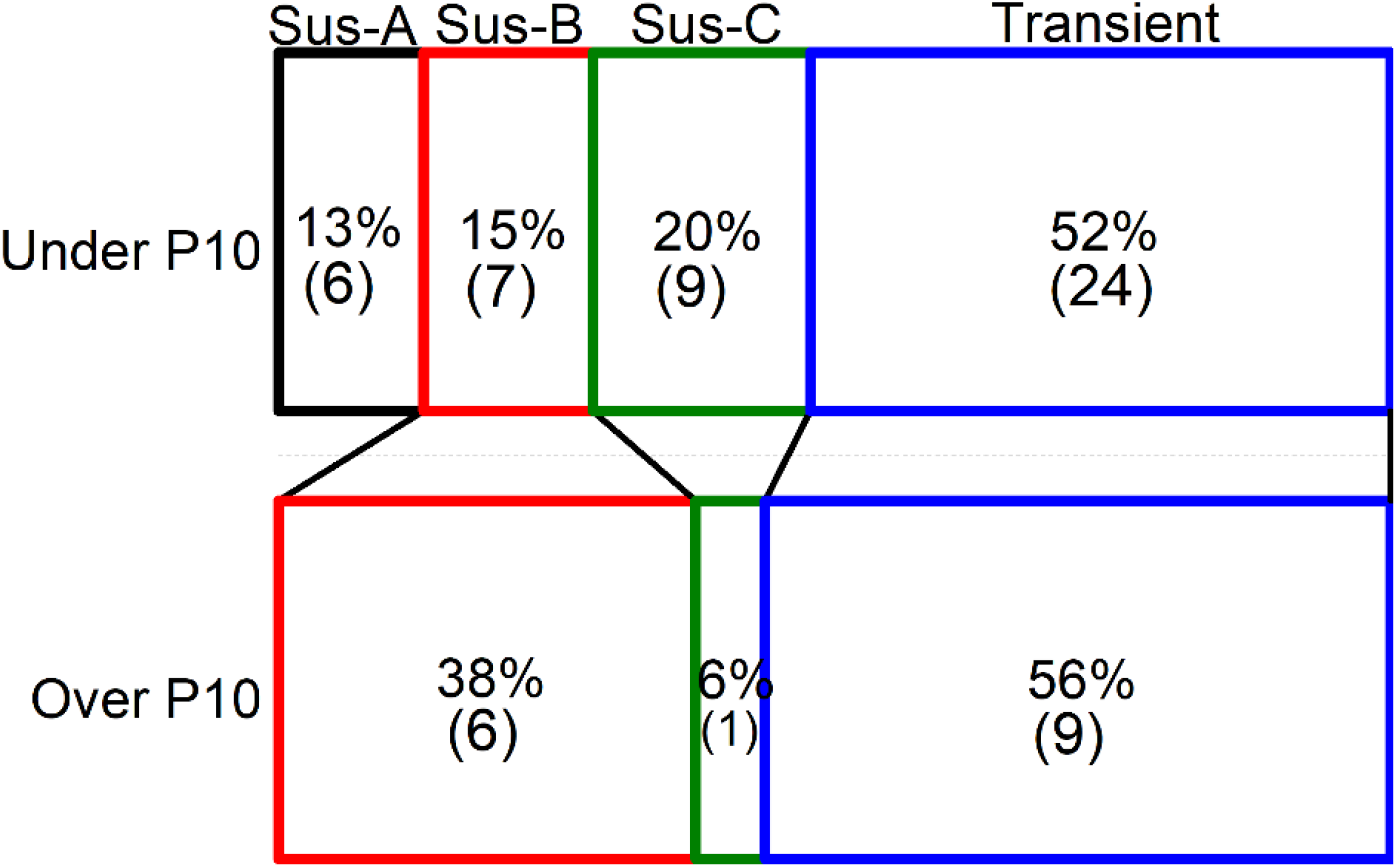
Changes in firing patterns and Na_V_ conductance density with age. Change in distribution of firing pattern in younger and older VGNs.

**Figure S4.**
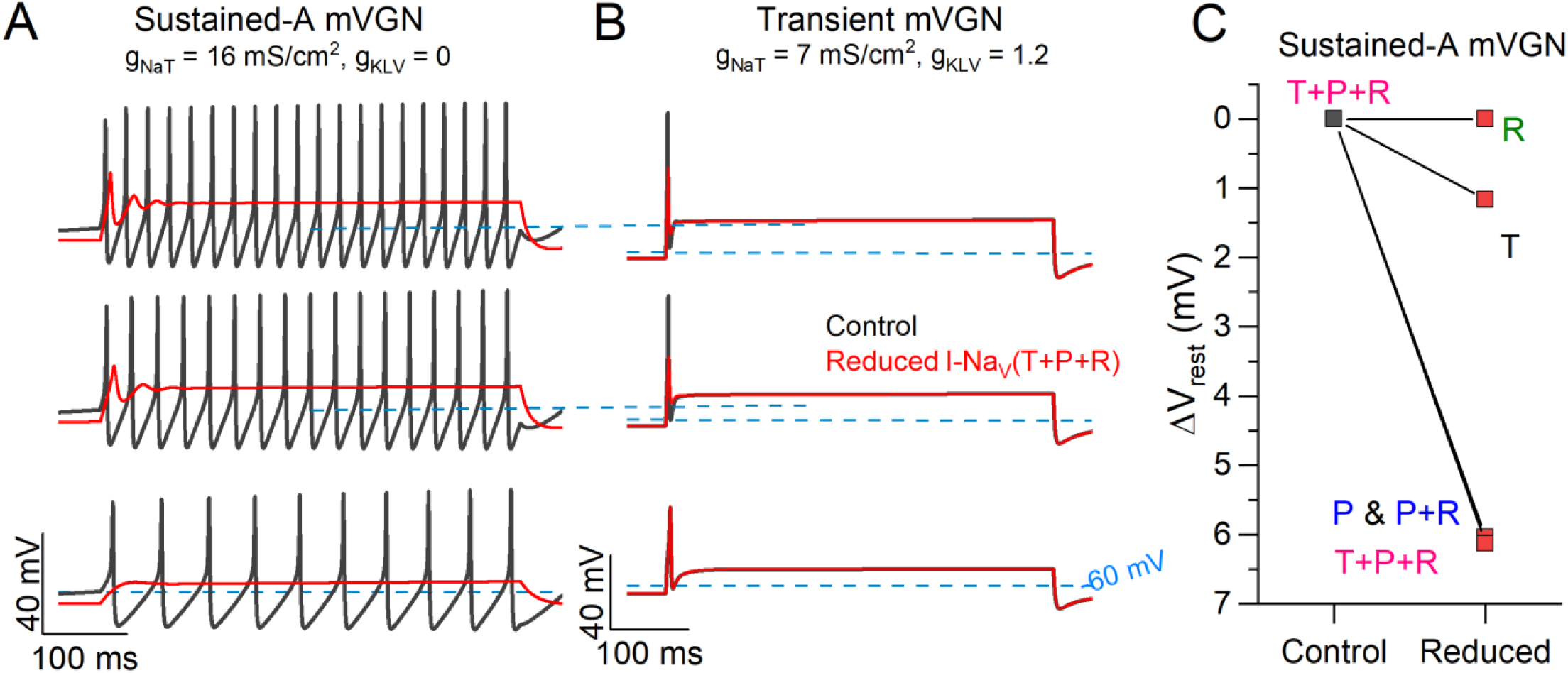
Reducing Na_V_ current modes replicates 4,9-ah-TTX block in model sustained and transient VGNs. **(A & B)** Firing patterns for a sustained-A (A) and transient (B) mVGNs. Reducing (red) I-Na_V_T by 70% and I-Na_V_R and I-Na_V_P by 90% replicates results observed with application of 4,9-ah-TTX in sustained-A mVGNs: increase in current threshold, reduction in spike height and spike rate, and slight hyperpolarization in V_rest_. **(B)** Transient mVGNs had reduced spike height and (not shown) greater current threshold. **(C)** Na_V_P and Na_V_T drive hyperpolarization in sustained mVGNs. Each component was reduced by the estimated 4,9-ah-TTX block: Na_V_R and Na_V_P by 90% and Na_V_T by 70%. Reducing (“blocking”) components in various combinations shows that blocking Na_V_T and Na_V_P hyperpolarizes resting membrane potential.

**Figure S5.**
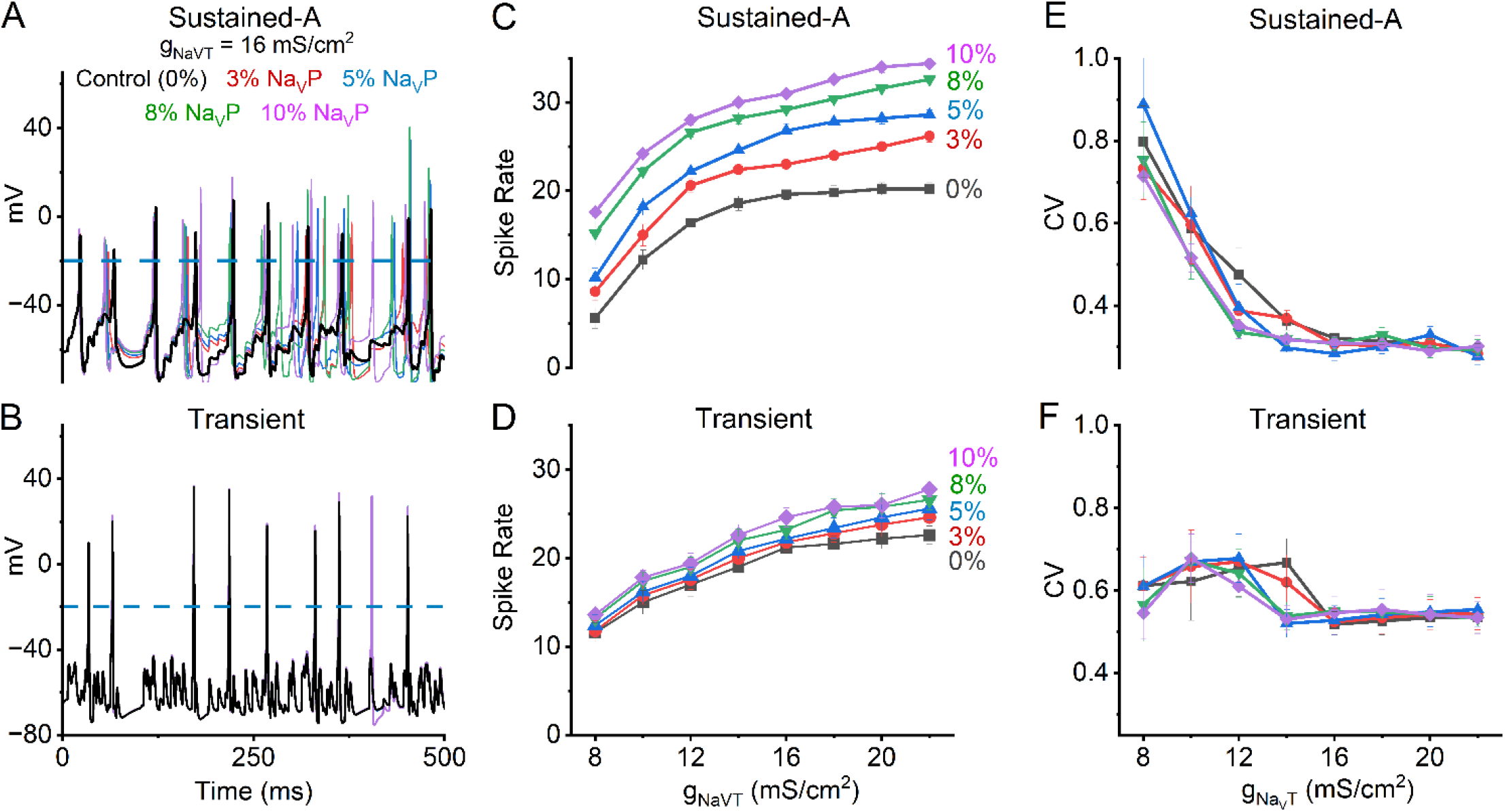
Increasing I-Na_V_P further increases spike rate. **(A & B)** Firing patterns for a sustained-A (A) and transient (B) mVGNs with I-Na_V_T density of 16 mS/cm^2^ with different amounts of I-Na_V_P: 0% (control, black), 3% (red), 5% (blue), 8% (green) and 10% (purple). **(C & D)** Summary of spike rates for sustained-A and transient mVGN under each level of I-Na_V_P. Increasing I-Na_V_P increased spike rate in both mVGNs. (**E & F)** A summary of spike train regularity as measured by CV. Increased I-Na_V_P had no effect on CV in either mVGN.

### 13.2 Supplementary Tables

**Table S1:**
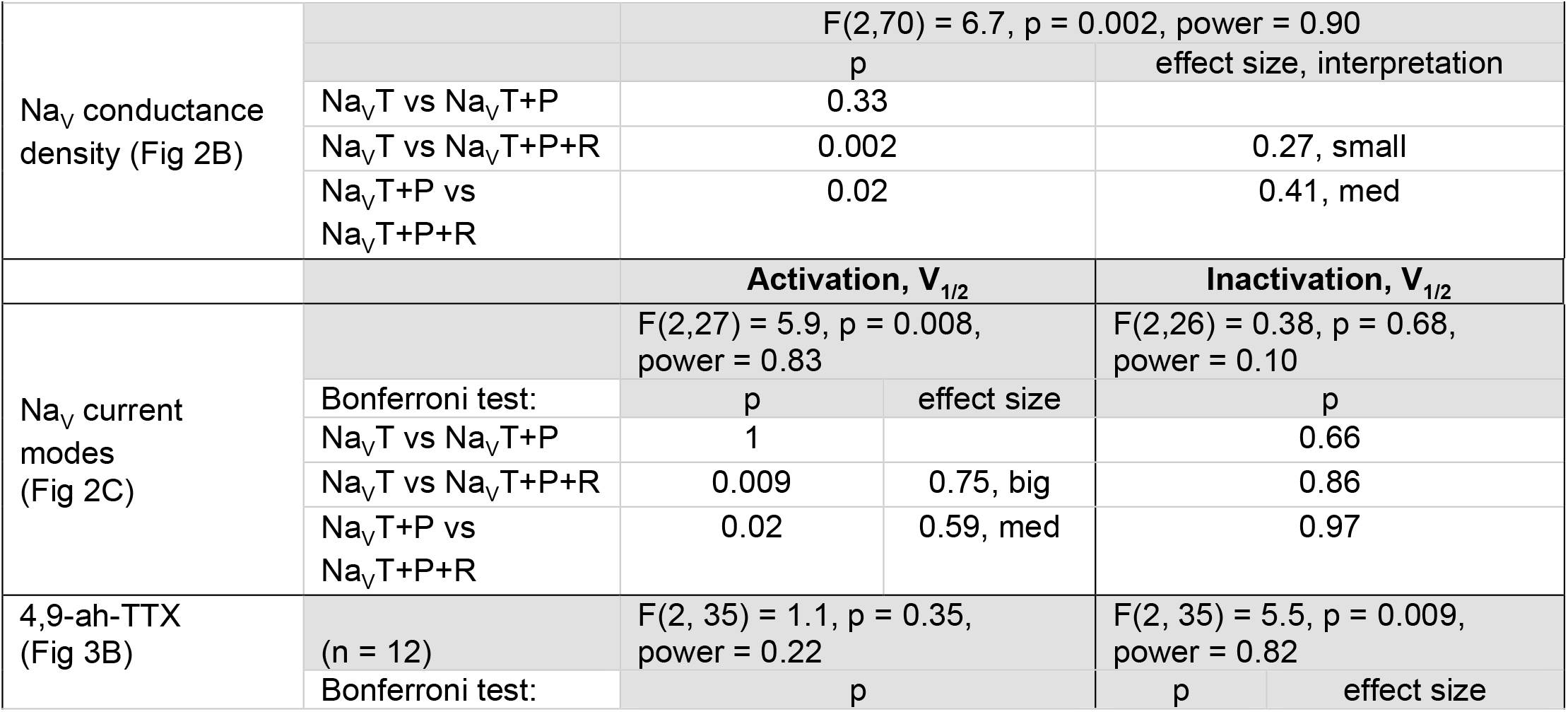
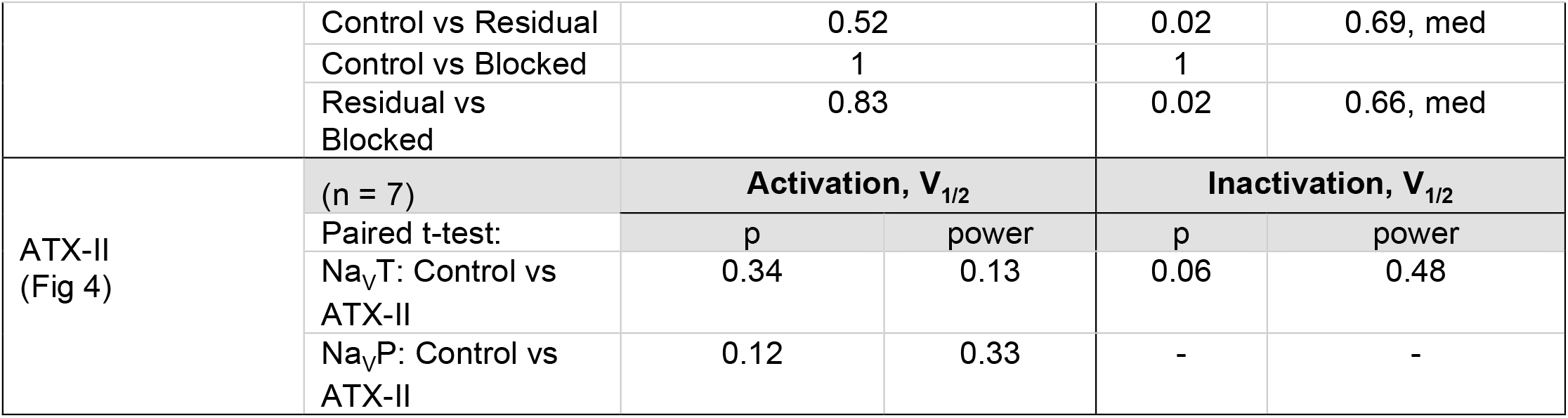
Summary of one-way ANOVA tests for voltage clamp experiments.

**Table S2:** Firing pattern passive properties (one-way 4-factor ANOVA)

| **Table S2: Firing pattern passive properties (one-way 4-factor ANOVA)** |  |  |  |  |
| --- | --- | --- | --- | --- |
| Firing pattern | Current threshold (pA)<br>(n) | $V_{rest}$ (mV) | $R_{in}$ (MΩ) | Cm (pF) |
| Sustained-A | 133.3 ± 16.7<br>(3) | -60.0 ± 2.2<br>(6) | 1020.0 ± 247.8<br>(5) | 21.9 ± 2.3<br>(6) |
| Sustained-B | 261.1 ± 26.1<br>(9) | -64.1 ± 1.8<br>(13) | 852.2 ± 280.6<br>(9) | 16.9 ± 1.8<br>(12) |
| Sustained-C | 295.5 ± 34.7<br>(11) | -63.5 ± 1.3<br>(10) | 475.3 ± 128.8<br>(7) | 15.1 ± 2.1<br>(9) |
| Transient | 426.8 ± 33.9<br>(28) | -65.7 ± 0.9<br>(33) | 464.9 ± 73.8<br>(20) | 16.1 ± 1.1<br>(31) |
| p | 0.001 | 0.11 | 0.08 | 0.18 |
| power | 0.94 | 0.51 | 0.56 | 0.41 |

**Table S3:** Direct comparisons of AP waveform differences (one-way 4-factor ANOVA)

|  |  |  |  |  |  |  |  |  |
| --- | --- | --- | --- | --- | --- | --- | --- | --- |
| | Current threshold (pA) | | Spike height ($V_{AP}$) (mV) | | Peak dV/dt (mV/ms) | | AHP (mV) | |
|  | p | Effect size | p | Effect size | p | Effect size | p | Effect size |
| Sustained-B vs Sustained-A | 0.35 |  | 0.08 |  | 0.79 |  | **0.009** | 0.6 |
| Sustained-C vs Sustained-A | 0.58 |  | 0.05 |  | 0.31 |  | **0.0005** | 0.9 |
| Sustained-C vs Sustained-B | 0.95 |  | 0.98 |  | 0.71 |  | 0.46 |  |
| Transient vs Sustained-A | **0.01** | 0.4 | **0.004** | **0.6** | **0.01** | 0.5 | **0.000006** | 1.0 |
| Transient vs Sustained-B | **0.03** | 0.4 | 0.66 |  | **0.03** | 0.3 | **0.04** | 0.5 |
| Transient vs Sustained-C | 0.08 |  | 0.92 |  | 0.58 |  | 0.77 |  |

**Table S4:** Pharmacological effects on AP waveform (paired t-test)

c
| **Table S4: Pharmacological effects on AP waveform (paired t-test)** |  |  |  |  |  |  |  |  |
| --- | --- | --- | --- | --- | --- | --- | --- | --- |
| **4,9-ah-TTX** | $V_{rest}$ (mV)<br>Transient | $V_{rest}$ (mV)<br>Sustained | Time-to-peak (ms) | Spike Width (ms) | AHP (mV) | $V_{AP}$ (mV) | Voltage threshold (mV) | Peak dV/dt (mV/ms) |
| Control (n = 12) | -59.5 ± 1.5 | -57.6 ± 3.3 | 4.1 ± 0.3 | 2.9 ± 0.5 | 0.0 ± 1.9 | 74.4 ± 6.4 | -42.3 ± 2.3 | 102 ± 20 |
| Drug | -63.6 ± 2.9 | -66.8 ± 3.4 | 4.8 ± 0.4 | 3.9 ± 0.6 | 6.8 ± 1.2 | 55.1 ± 7.0 | -43.3 ± 3.2 | 64.4 ± 20 |
| p | 0.16 | **0.03** | **0.05** | 0.13 | **0.006** | **0.01** | 0.90 | **0.03** |
| Effect size |  | 0.67 | 0.42 | 0.19 | 0.72 | 0.49 | 0.05 | 0.41 |
| Power | 0.27 | 0.67 | 0.53 | 0.31 | 0.86 | 0.76 | 0.05 | 0.62 |
| **ATX-II** | $V_{rest}$ (mV) | | | | | | | |
| Control (n = 7) | -59.2 ± 0.9 |  | 3.8 ± 0.3 | 1.2 ± 0.1 | 4.8 ± 0.8 | 92.7 ± 5.9 | -49.7 ± 2.1 | 229 ± 30 |
| Drug | -64.0 ± 2.2 |  | 5.5 ± 0.8 | 2.4 ± 0.6 | 4.1 ± 1.5 | 92.8 ± 6.3 | -52.2 ± 3.5 | 209 ± 50 |
| p | 0.07 |  | 0.07 | 0.09 | 0.58 | 0.97 | 0.60 | 0.41 |
| Effect size | 0.83 |  | 0.45 | 0.35 | 0.10 | 0.01 | 0.23 | 0.14 |
| Power | 0.45 |  | 0.45 | 0.41 | 0.07 | 0.05 | 0.07 | 0.11 |

**Table S5:** AP waveform differences between model VGNs.

| **Table S5: AP waveform differences between model VGNs** |  |  |  |  |  |  |  |  |  |  |  |
| --- | --- | --- | --- | --- | --- | --- | --- | --- | --- | --- | --- |
| mVGN<br>(Figure 8) |  |  | 1<sup>st</sup> AP: |  |  |  |  | ISI<br>(ms) | 2<sup>nd</sup> AP: |  |  |
|  | Nav<br>modes | Vrest<br>(mV) | Time-to-peak<br>(ms) | V<sub>AP</sub><br>(mV) | Spike<br>width<br>(ms) | Peak<br>dV/dt<br>(mV/ms) | AHP<br>(mV) |  | Time-to-peak<br>(ms) | V<sub>AP</sub><br>(mV) | Spike<br>width<br>(ms) |
| Sustained-A | T | -59.6 | 2.6 | 96 | 1 | 143.4 | -6.9 | 8.6 | 6.4 | 69.2 | 1.4 |
|  | T+P | -54.7 | 2.3 | 77.9 | 1.2 | 100.6 | -9.6 | 8.3 | 6 | 61.8 | 1.4 |
|  | T+R | -59.6 | 2.6 | 96 | 1 | 143.4 | -3.3 | 7.2 | 5.1 | 63.7 | 1.6 |
|  | T+P+R | -54.7 | 2.3 | 77.9 | 1.2 | 100.6 | -9.4 | 8.2 | 5.9 | 61.8 | 1.5 |
|  | T+ | -59.6 | 2.5 | 101.6 | 0.9 | 162.4 | -7.3 | 8.6 | 6.2 | 74 | 1.3 |
| Sustained-B | T | -61.7 | 3.9 | 85.9 | 1.2 | 105.1 | -5.6 | 12.9 | 10.3 | 57 | 2.3 |
|  | T+P | -61.1 | 3.8 | 84.7 | 1.2 | 106 | -6.1 | 12.4 | 9.8 | 58.8 | 1.8 |
|  | T+R | -61.7 | 3.9 | 85.9 | 1.2 | 105.1 | -5.4 | 12.8 | 10.2 | 57.1 | 2.3 |
|  | T+P+R | -61.1 | 3.8 | 84.7 | 1.1 | 106 | -6 | 12.3 | 9.7 | 58.9 | 2 |
|  | T+ | -62.7 | 3.8 | 92.7 | 1.1 | 123.1 | -5.3 | 12.7 | 10.2 | 63.7 | 1.9 |
| Sustained-C | T | -65.2 | 5.1 | 86.6 | 1.4 | 92.8 | -0.8 | 16.6 | 13.5 | 48.5 | 3.4 |
|  | T+P | -64.7 | 4.9 | 86.5 | 1.5 | 94.2 | -1.2 | 15.9 | 12.7 | 51.2 | 3.2 |
|  | T+R | -65.2 | 5.1 | 86.6 | 1.4 | 92.8 | -0.7 | 16.5 | 13.4 | 48.8 | 3.8 |
|  | T+P+R | -64.7 | 4.9 | 86.4 | 1.5 | 94.2 | -1.2 | 15.7 | 12.1 | 51.5 | 3.2 |
|  | T+ | -65.2 | 5 | 91.9 | 1.5 | 108.7 | -1.5 | 16.1 | 12.9 | 53.7 | 2.8 |
| Transient | T | -64.3 | 2.5 | 77.6 | 1.5 | 77.9 | 5.4 |  |  |  |  |
|  | T+P | -64.2 | 2.5 | 77.9 | 1.5 | 79.7 | 5.3 |  |  |  |  |
|  | T+R | -64.2 | 2.5 | 77.9 | 1.5 | 77.9 | 5.3 |  |  |  |  |
|  | T+P+R | -64.2 | 2.5 | 77.9 | 1.5 | 79.7 | 5.3 |  |  |  |  |
|  | T+ | -64.3 | 2.5 | 82 | 1.4 | 90.9 | 4.7 |  |  |  |  |

**Table S6:**
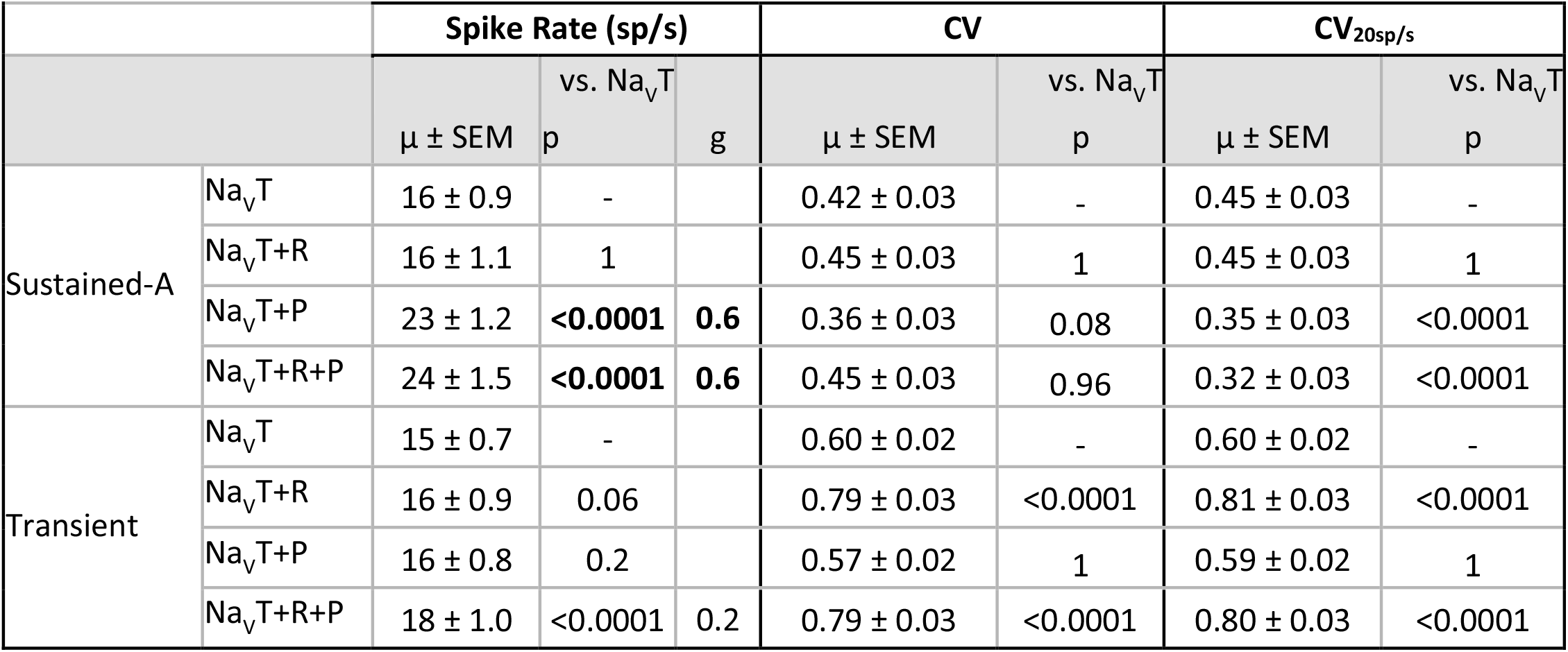
Effects of I-Na_V_P and I-Na_V_R on spike rate and CV in model VGNs (two-way 4-factor ANOVA)

