## Supplementary figures and images for "Effects of transient, persistent, and resurgent sodium currents on excitability and spike regularity in vestibular ganglion neurons"

### Supplemental Fig 1

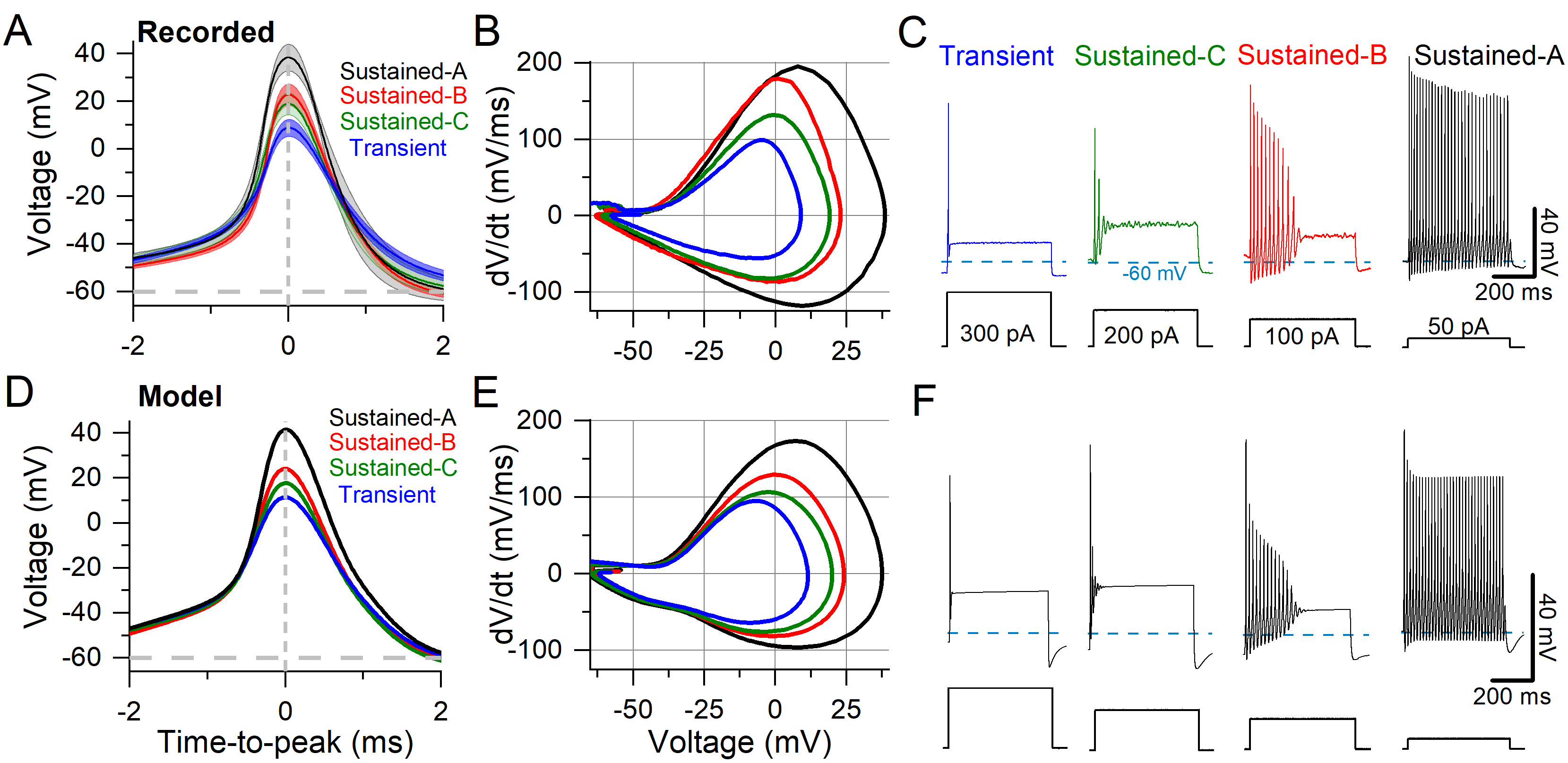

### Supplemental Fig 2

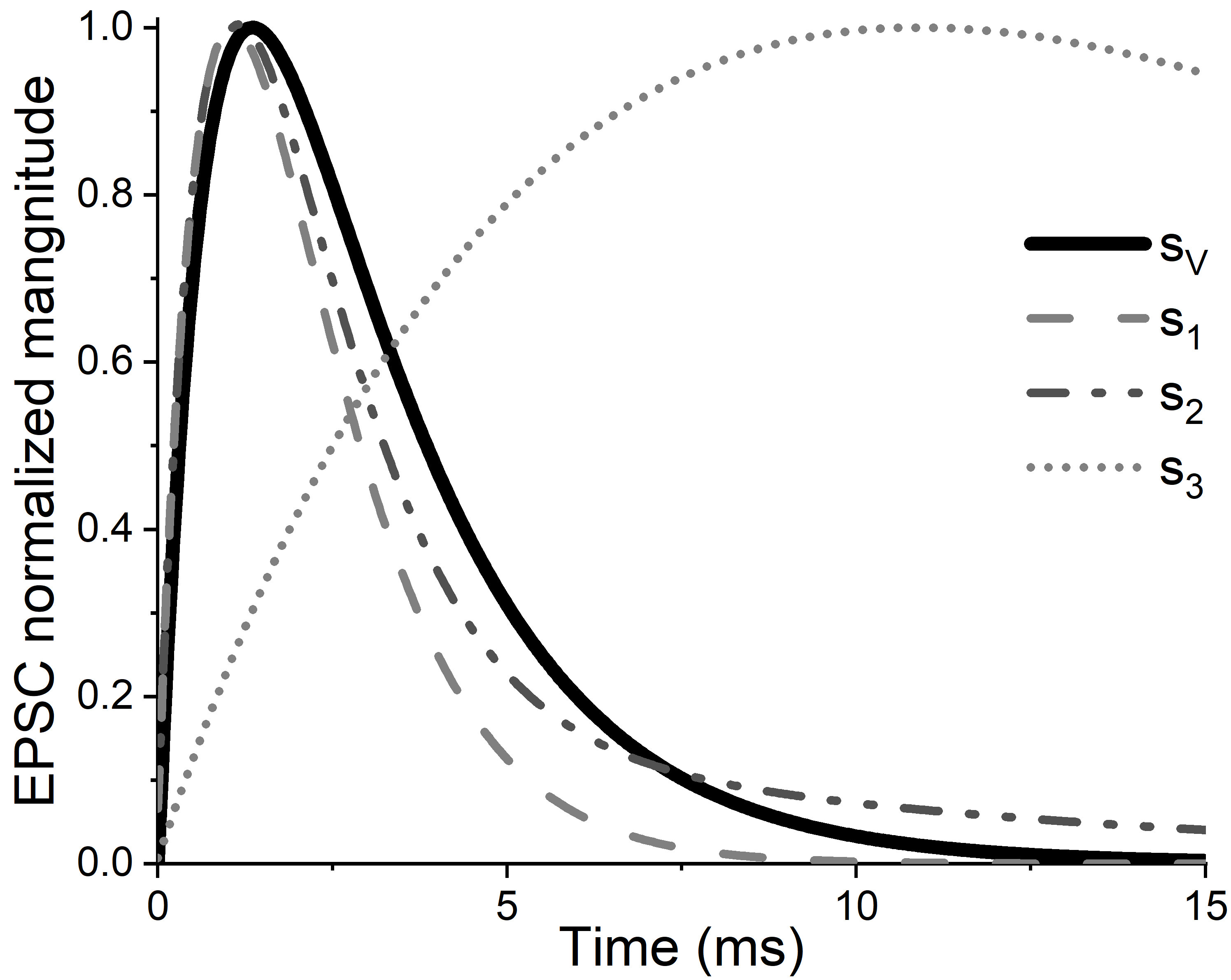

### Supplemental Fig 3

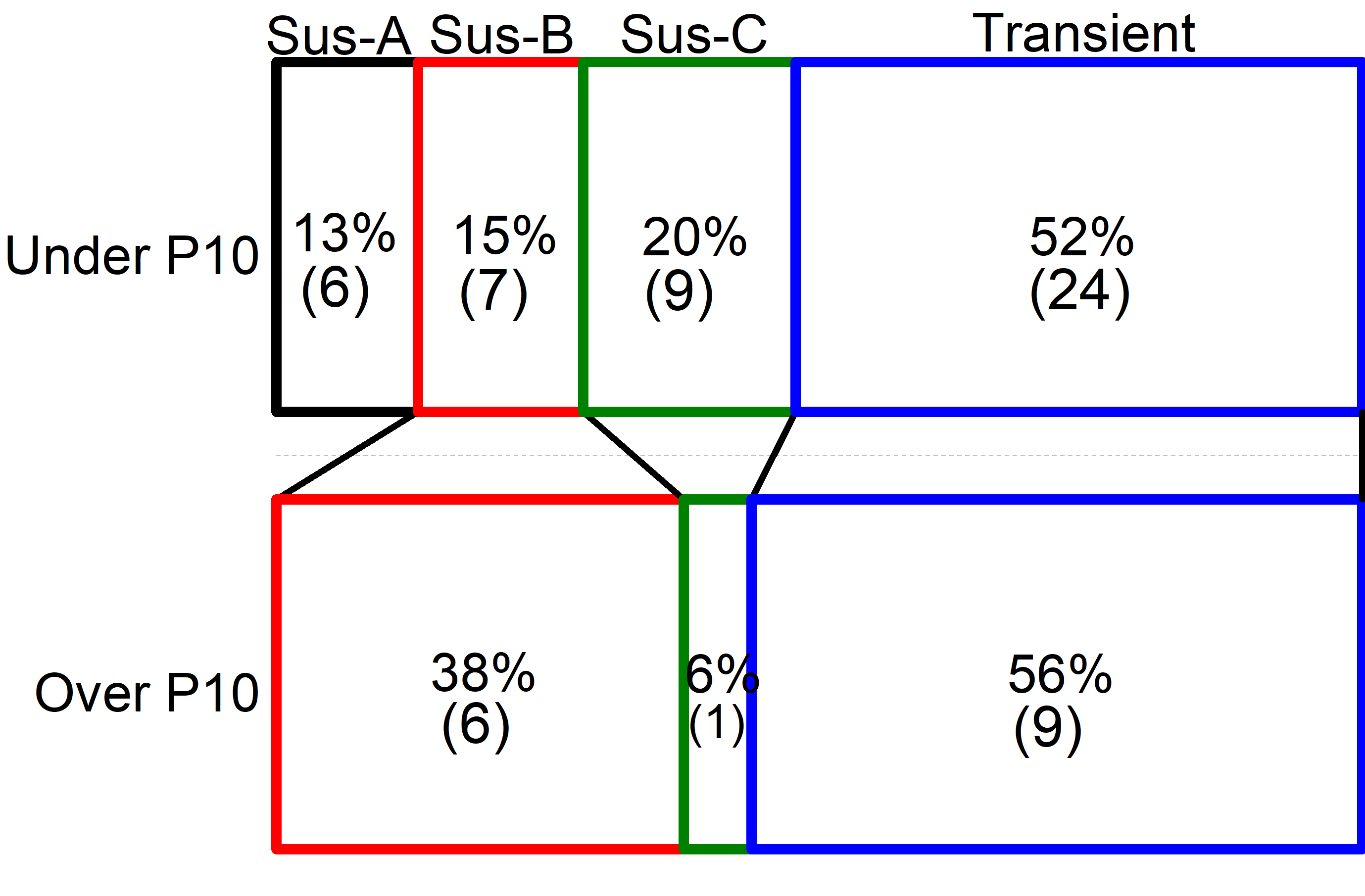

### Supplemental Fig 4

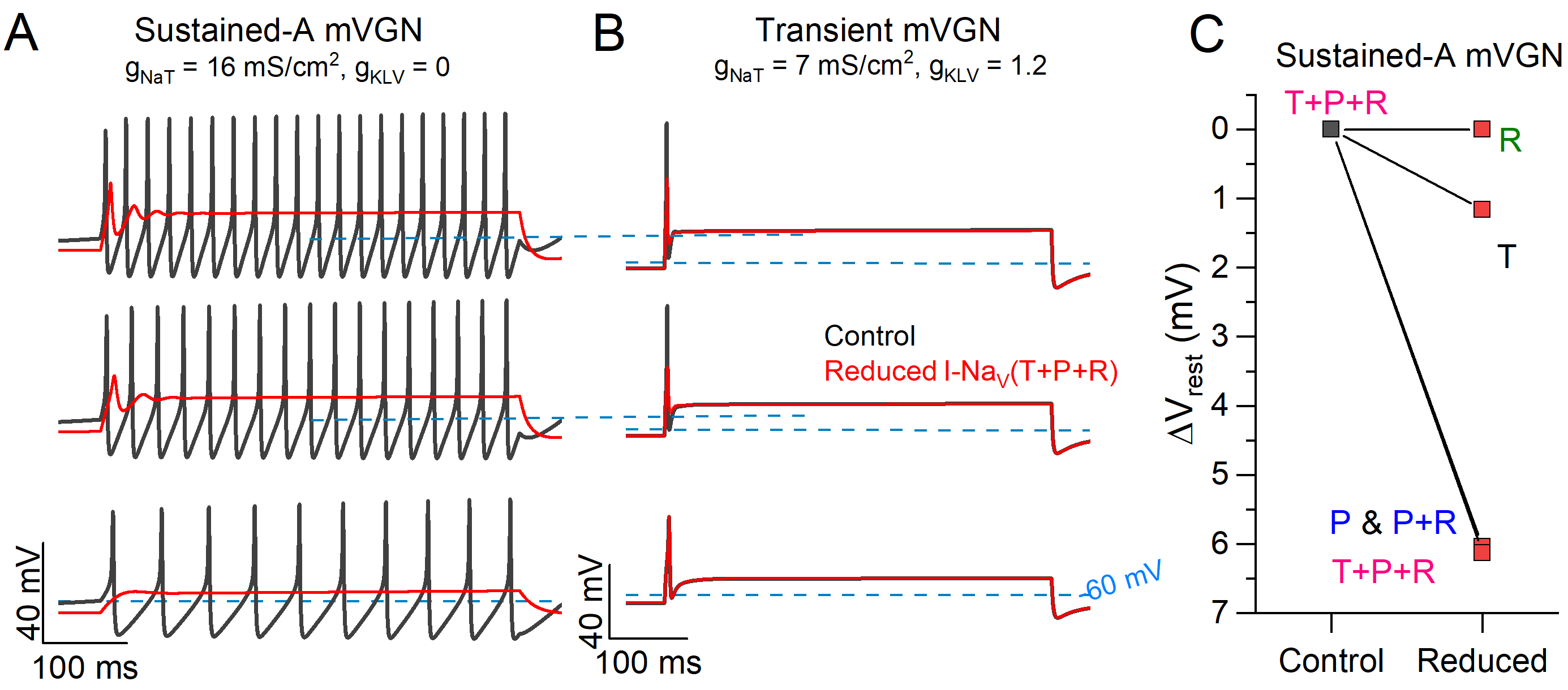

### Supplemental Fig 5

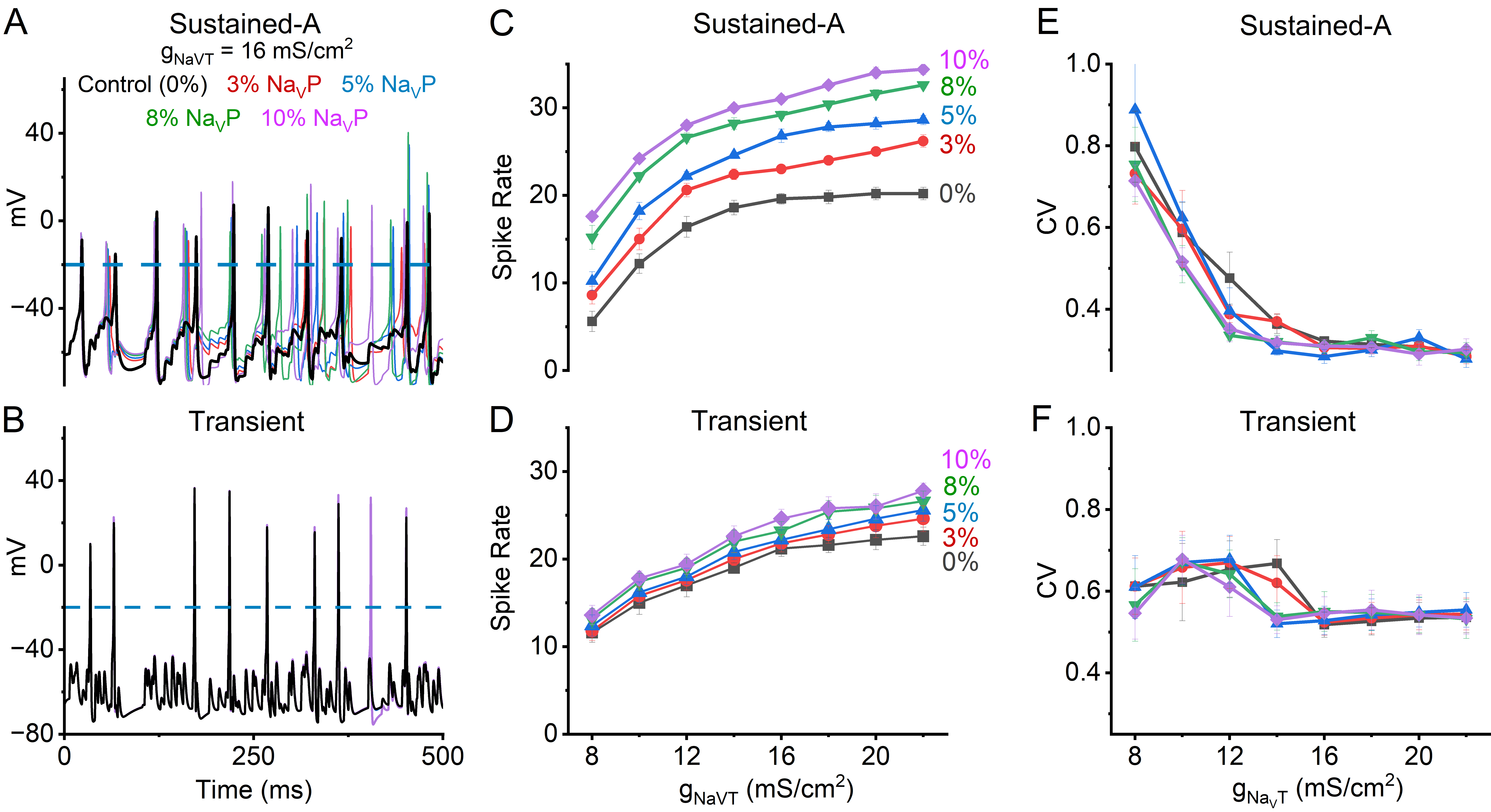
